# HIV-1 infection induces Vif-mediated SUMOylation of host RNA splicing factors important for proper viral RNA splicing

**DOI:** 10.1101/2025.03.26.645526

**Authors:** Demetra P. Kelenis, Jeffrey R. Johnson, Simone Sidoli, Ann Emery, Ronald Swanstrom, Luke J. Hawkins, Kaila Mckie, Tegh Pawar, Stephen P. Goff

## Abstract

HIV-1 exploits host cell post-translation modifications (PTMs) to facilitate production of infectious particles. These modifications include SUMOylation, a dynamically regulated PTM involving covalent attachment of small ubiquitin-like modifiers (SUMOs) to lysine (K) residues of target proteins. SUMOylation modulates the activity of thousands of proteins and multiple fundamental host cellular processes, including pathways hijacked by HIV-1 to promote infection and spread. The SUMOylation of several proteins during HIV-1 infection has been characterized. However, the broad effects of HIV-1 infection on the SUMOylation of the host cell proteome is largely unknown. To date, SUMOylation has not been explored by large-scale proteomics in the context of HIV infection, where many SUMO-regulated host dependency factors remain to be identified. In this study, we performed a proteome-wide, mass spectrometry (MS)-based screen to identify proteins that are SUMOylated during HIV-1 infection. Here, and in biochemical assays, infection with HIV-1 led to the widespread increased SUMOylation of the heterogeneous nuclear ribonucleoprotein (HNRNP) A/B proteins, a protein family central to the regulation of alternative splicing. Intriguingly, this phenotype was found to be driven by expression of the HIV-1 Viral Infectivity Factor (Vif), suggesting a novel function for this protein aside from APOBEC3G degradation. We selected HNRNPA2B1 (A2/B1) and HNRNPA3 for further study, where depletion of these proteins led to the altered splicing of HIV-1 viral RNAs and dramatically reduced HIV-1 infectivity. Considering the enrichment of SUMOylation sites within the RNA-binding domains of the HNRNPA/B family, our data suggest a novel mechanism involving HIV-1-induced, Vif-mediated SUMOylation of host RNA splicing factors as a means to regulate HIV-1 alternative splicing. Broadly, our findings suggest that infection with HIV-1 alters the SUMOylation of many unexplored host cellular proteins, and provides a significant proteomic resource for their future mechanistic study.

## INTRODUCTION

SUMOylation is a reversible, dynamic post-translational modification (PTM) involving covalent attachment of Small Ubiquitin-like Modifier (SUMO) proteins to lysine (K) residues of substrate proteins. Like ubiquitination, SUMOylation occurs via a three-step, ATP-dependent enzymatic cascade including an E1 activating enzyme (SAE1/SAE2), E2 conjugase (UBC9), and for some substrates, an E3 ligase.^1,2,3^ Unlike ubiquitination, SUMOylation does not typically target proteins for degradation and instead influences protein activity, often by promoting non-covalent interactions with SUMO Interacting Motif (SIM)-containing cofactors. Conserved from yeast to mammals, SUMOylation modulates the activity of thousands of proteins and multiple fundamental cellular process such as nucleocytoplasmic transport, intracellular trafficking, signal transduction, cell cycle control, and DNA and RNA metabolism.^4,5,6,7^ SUMOylation also provides cells with a rapid, versatile mechanism to alter proteomic activity in response to various stimuli, including viral infection. In turn, both DNA and RNA viruses have been shown to manipulate host cell SUMOylation by directly targeting the SUMO pathway.^6,2,3,8^

Like other viruses, human immunodeficiency virus type 1 (HIV-1) is entirely dependent upon the host cell machinery for completion of the viral life cycle and is known to manipulate this machinery to ensure infection and spread. In part, HIV-1 accomplishes this by utilizing a variety of post-translational modifications (PTMs) to “re-wire” essential host cellular processes, enabling these pathways to support HIV-1 replication. HIV-1 directly exploits multiple PTMs, including glycosylation, myristoylation, phosphorylation, ubiquitination, and SUMOylation.^9,10,11^ HIV-1 has a special dependence on alternative mRNA splicing, a process regulated by the SUMOylation of splicing factors in uninfected cells.^12,13^ However, despite the dependence of HIV-1 on SUMO-regulated host cell pathways, the exploitation of SUMOylation as a means to remodel these pathways during HIV-1 infection is largely unexplored.

The study of SUMOylation during HIV-1 infection has primarily relied on targeted analysis of HIV-1 viral proteins and host cellular proteins already known to mediate HIV-1 replication. For instance, SUMOylation of the HIV-1 viral protein p6 has been shown to compete with ubiquitination at the same residue, inhibiting recruitment of ESCRT machinery during virus production, while SUMOylation of HIV-1 integrase influences the integration efficiency of the HIV-1 DNA into the host cell genome.^14,15,16,17,15^ SUMOylation of unintegrated chromatinized HIV-1 DNA has been shown to mediate the establishment of HIV-1 latency and a recent study suggests that SUMOylation may also regulate proteins involved in HIV-1 viral latency maintenance in microglia.^18,19^ The SUMOylation of several HIV-1 host factors, such as NFKBIA, LEDGF/p75 (PSIP1), SAMHD1, and CDK9 have also been shown to influence HIV-1 replication,^9,20,21,22,23,24,25^ or latency.^26^ SUMO-SIM interactions are also involved in TRIM5α-mediated retroviral restriction.^27^ To date however, SUMOylation had not been studied by system-wide proteomics in the context of HIV-1 infection, where many SUMO-regulated host dependency factors remain to be identified and the functional significance of these modifications explored.

In this study, we performed an unbiased, proteome-wide mass spectrometry (MS)-based screen to identify host cell proteins that are SUMOylated in response to HIV-1 infection in human cells. We experimentally tested the effects of HIV-1 infection on the SUMOylation of 30 proteins identified by MS. Infection induced the increased SUMOylation of the heterogeneous nuclear ribonucleoprotein (HNRNP) A/B family of splicing factors. We further found that their SUMOylation is driven by expression of the HIV-1 viral protein Vif. We focused our attention on HNRNPA2B1 (A2/B1) and HNRNPA3, two HNRNPA/B family members that were heavily SUMOylated upon HIV-1 infection across multiple cell line models. Quantification of HIV alternative splicing following HNRNPA2B1 and HNRNPA3 depletion combined with computational analysis of their predicted SUMOylation sites suggest a potential role for this modification in mediating the splicing of HIV-1 viral RNAs.

Together, our findings suggest a mechanism by which HIV-1 infection induces Vif-mediated SUMOylation of the HNRNPA/B family, a protein family with a central role in host cellular and HIV-1 viral RNA metabolism. These findings provide novel insight into the regulation of HIV-1 alternative splicing and uncover a previously unappreciated function for the viral protein Vif.

## RESULTS

### Proteome-wide screen identifies HIV-1-induced SUMO substrates

The human genome encodes five SUMO paralogs (SUMO1-5). SUMO1-3 are expressed ubiquitously, while the expression of SUMO4 and 5 is restricted to specific tissues. SUMO2 and 3 are 97% identical in amino acid sequence and are often grouped together as SUMO2/3. SUMO1 and SUMO2/3 share only 47% sequence homology and are regarded as functionally distinct.^1,2,3^ To investigate the dynamics of SUMOylation in response HIV-1 infection by the main SUMO paralogs, we utilized a previously described proteomic strategy^3,28^ to identify host cell proteins that change in SUMO1 or SUMO2 modification status following infection with HIV-1.

We generated stable cell lines expressing SUMO1 or SUMO2 fused to an N-terminal 3xFLAG tag. To facilitate mapping by MS, our SUMO constructs also included previously described C-terminal glutamine to arginine (Q to R) mutations (3xFLAG-SUMO1^Q92R^, 3xFLAG-SUMO2^Q88R^). These mutations introduce a trypsin cleavage site near the C-terminus of SUMO, leaving a unique five amino acid remnant on the SUMO1 versus SUMO2-modified peptides following tryptic digest (Figure 1A).^3,28^ 3xFLAG-tagged SUMO1 and SUMO2 mutants were expressed at levels comparable to endogenous WT SUMOs, and were conjugated to target proteins in the respective cell lines (Supplemental Figure 1A). To enable comparison between different strategies for enriching SUMOylated substrates, we also generated cell lines stably expressing HIS_6_-tagged SUMO1^Q92R^ or SUMO2^Q88R^ (Figure 1A). Both 3xFLAG and HIS_6_-tagged SUMO1^Q92R^ immunoprecipitated with RANGAP1, a well-characterized SUMO1 substrate (Supplemental Figure 1B).^29^ This suggests that the combinations of N-terminal tags and C-terminal mutations utilized did not interfere with the conjugation of SUMO mutants to endogenous target proteins. HIV-1 infection efficiency and reporter expression was also unaffected in the 3xFLAG or HIS_6_-tagged SUMO cell lines, indicating that expression of exogenous SUMOs did not interfere with HIV-1 infection (Supplemental Figure 1C).

**Figure 1.**
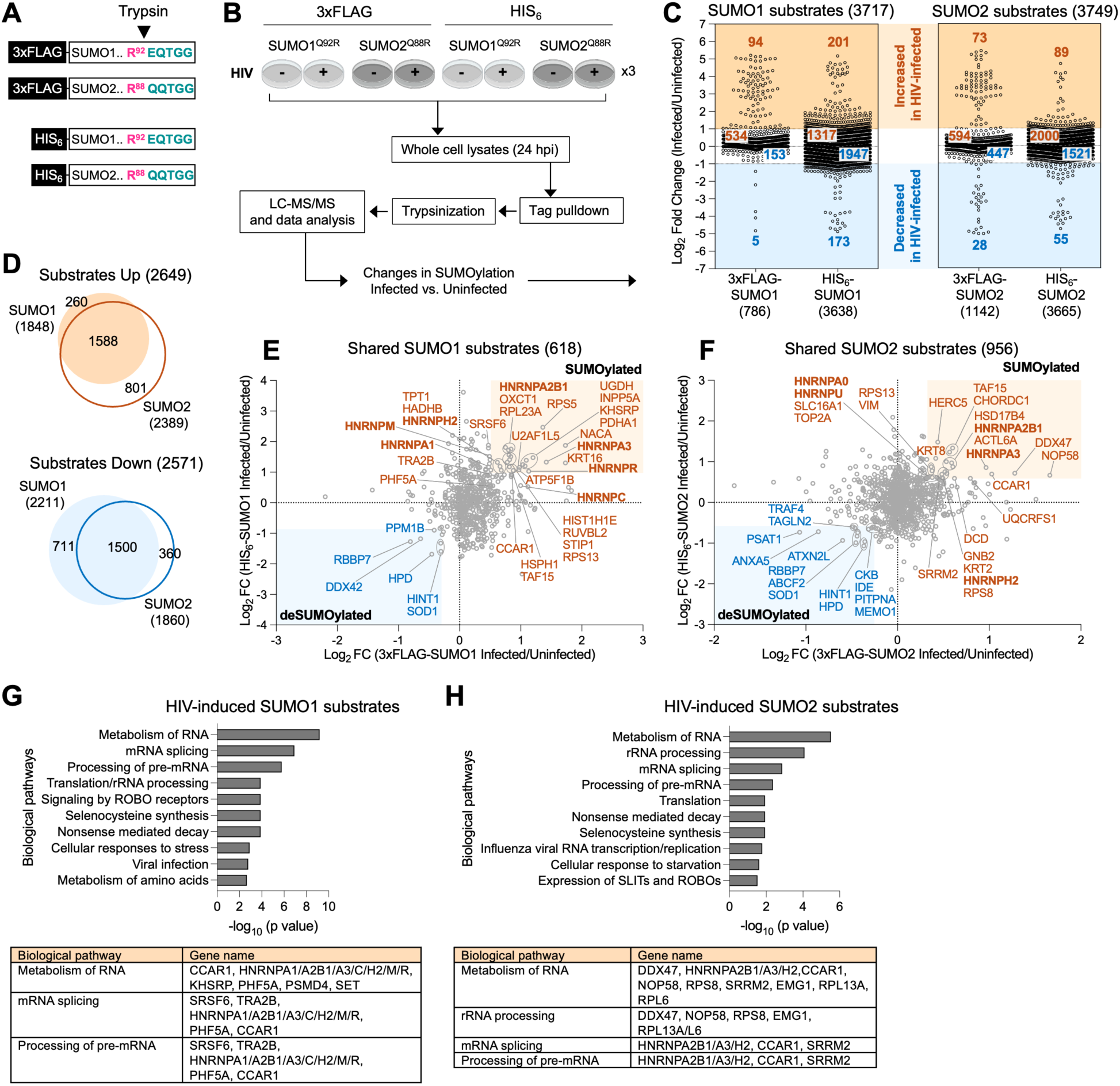
Proteome-wide identification of proteins SUMOylated during HIV-1 infection. **(A)** 3xFLAG and HIS_6_-tagged mutant SUMO expression vectors. Tryptic digest leaves an -EQTGG or -QQTGG signature remnant on SUMO1 or SUMO2 modified peptides, respectively. **(B)** Overview of proteomic strategy to identify proteins with a change in SUMOylation status during HIV-1 infection. Three replicates were performed per condition. **(C)** Log_2_ fold changes in SUMO1 and SUMO2 substrate abundance (y-axis, Infected/Uninfected) in the 3xFLAG or HIS_6_-tagged SUMO cell lines following HIV-1 infection (x-axis). Orange and blue shaded areas designate substrates with a Log_2_ fold change >1 or < -1 (>2-fold linear change up or down, respectively). The numbers of substrates identified in each cell line and each category are indicated. **(D)** Overlap between SUMO1 and SUMO2 substrates that were increased or decreased by HIV-1 infection. **(E, F)** Graphs indicating changes in abundance to SUMO1 (E) and SUMO2 (F) substrates derived from 3xFLAG-tagged SUMO cells (x-axis) versus the respective HIS_6_-tagged SUMO cells (y-axis) in response to HIV-1 infection. Log_2_FC = Log_2_ fold change. Substrates with fold changes in the top or bottom quartiles in both the 3xFLAG- and HIS_6_-tagged SUMO samples are highlighted, in addition select splicing factors indicated in G, H. **(G, H)** Significantly enriched pathways in the HIV-induced SUMO1 and SUMO2 substrates common to the 3xFLAG- and HIS_6_-tagged SUMO samples shown in parts E and F. Select genes included in these pathways are shown in the tables below.

The 3xFLAG- and HIS_6_-tagged SUMO1 and SUMO2 cell lines were used to identify host proteins that change in SUMOylation status in response to HIV-1 infection. Cells were either left uninfected, or infected with single-cycle HIV-1 (pNL4-3 deltaENV-EGFP) at an infection efficiency of ∼60% and harvested at 24 hours post-infection (hpi). 3xFLAG-SUMO-conjugated proteins were isolated from cell lysates using an anti-FLAG antibody, while HIS_6_-SUMO-conjugated proteins were enriched using a Nickel-NTA column. SUMO-conjugated proteins were isolated under denaturing conditions to dissociate non-covalent interactors and inactivate SUMO proteases. Proteins were trypsinized and analyzed by liquid chromatography–tandem mass spectrometry (LC-MS/MS) for protein identification and abundance quantification (Figure 1B).

This approach identified 3,717 total SUMO1-modified proteins and 3,749 SUMO2-modified proteins (Supplemental Table 1, Tabs A and B), in addition to a small number of -EQTGG (SUMO1) or -QQTGG (SUMO2)-modified peptides (Supplemental Table 2). Among these proteins, there were many that showed increased or decreased levels of SUMOylation in the HIV-1-infected samples relative to the uninfected controls. We first compared the total list of SUMO-modified substrates identified in this study to a database of SUMOylated proteins comprised of data from 22 proteomic studies.^30^ Here, ∼54% of all SUMOylated proteins identified in our study overlapped with those previously reported in the database (Supplemental Figure 2A). Our data also included many well-known SUMO substrates such as RANGAP1, TOP2A, PML, SART1, SAFB2, RANBP2, PCNA, TRIM28, TRIM33, BLM, and BRCA1, as well as proteins whose SUMOylation status has been shown to modulate acute HIV-1 infection (NFKBIA, PSIP1, SAMHD1) or HIV-1 latency (PLK1, CDK9) (Supplemental Table 1, Tabs A and B).^9,20,21,22,23,31,26^ Together, these findings support the validity of our approach for identifying confirmed SUMO substrates, and suggest that there are many unexplored proteins that may become SUMO1 or SUMO2 targets upon HIV-1 infection.

The effect of HIV-1 infection on all SUMO1- or SUMO2-modified substrates identified in the 3xFLAG- or HIS_6_-tagged cell lines is shown in Figure 1C. The y-axis indicates the Log2 fold change in abundance of these proteins in the HIV-1-infected samples relative to the uninfected controls, and the x-axis indicates the respective 3xFLAG- or HIS_6_-tagged condition. On average, a slightly higher percentage of SUMO substrates were increased by HIV-1 infection (SUMOylated, 60%) versus decreased (deSUMOylated, 40%). For most of these proteins, infection with HIV-1 did not lead to a Log_2_ dynamic range larger than 1 (2-fold linear change up or down). Still, 463 SUMO1 substrates and 244 SUMO2 substrates increased or decreased more than 2-fold with HIV-1 infection, and some dramatically changed (Figure 1C) (Supplemental Table 1, Tabs C-F). Together, these findings suggest that HIV-1 infection leads to concomitant increase or decrease in SUMOylation of different sets of host proteins, with HIV-1 substantially altering the SUMOylation status or abundance of many substrates. Comparison between SUMO1 versus SUMO2 substrates that were increased or decreased following HIV-1 infection, respectively, revealed a high degree of overlap (Figure 1D) (Supplemental Table 1, Tab G), suggesting that most of the SUMO substrates we identified can be modified by either SUMO1 or SUMO2 during HIV-1 infection.

### HIV-1 infection leads to SUMOylation of HNRNPs and other splicing factors

To identify proteins that are consistently SUMOylated upon HIV-1 infection regardless of tag or enrichment strategy, we looked for substrates that had similar quantitative changes in SUMO1 or SUMO2 modification in response to HIV-1 infection in both the 3xFLAG-tagged and HIS_6_-tagged SUMO cell lines (Figure 1E, F). Among the SUMO1 substrates identified in both the 3xFLAG- and HIS_6_-tagged cell lines, there were numerous heterogeneous nuclear ribonucleoprotein (HNRNP) family members that were increased following infection with HIV-1, including HNRNPs A2B1, A3, H1, H2, C, F, K, M, R, U, A0 and A1 (Figure 1E) (Supplemental Table 1, Tab H). Similarly, among the SUMO2 substrates identified in both the 3xFLAG- and HIS_6_-tagged cell lines, HNRNPs A2B1, A3, A0, U, H2, H1, C, K, R, U, A1, and AB were enriched in the HIV-1-infected cells relative to the uninfected controls (Figure 1F) (Supplemental Table 1, Tab H). The shared SUMOylation of many these HNRNPs by both SUMO1 and SUMO2 upon HIV-1 infection is consistent with the high level of overlap in SUMO1 and SUMO2 substrates observed in our data (Figure 1D). Consistent with the central role of HNRNPs in regulating multiple aspects of RNA metabolism, including alternative splicing, the HIV-1-induced SUMO1 and SUMO2 substrates were significantly enriched for Reactome pathways “Metabolism of RNA”, “mRNA splicing”, and “Processing of capped intron-containing pre-mRNA”, among other RNA-related pathways (Figure 1G, H) (Supplemental Table 1, Tab I).^32^ The HIV-1-induced SUMO substrates also included additional splicing factors such as SRSF6, TRA2B, PHF5A, CCAR1, and SRRM2. (Figure 1G, H). Importantly, western blot analysis of 30 of these SUMO substrates revealed no substantial difference in expression level of the unmodified form of the substrate protein between HIV-1-infected versus uninfected cells, for all but 6 proteins (Supplemental Figure 2B). This suggests that a large proportion of the substrates we identified show specific increases in SUMO modification upon HIV-1 infection that cannot be explained by increased total protein abundance. All together, our mass spectrometry data indicates that infection with HIV-1 leads to a broad pattern of SUMOylation of HNRNPs and other mediators of alternative splicing.

### Experimental validation of HIV-1-induced SUMO targets

To confirm the HIV-1-induced SUMOylations detected by mass spectrometry, we selected 20 putative SUMO1 targets and 14 putative SUMO2 targets for experimental validation in immunoprecipitation (IP) assays (Figure 2A). In many prior studies, experimental validation of SUMO targets relied on overexpression of tagged SUMOs and/or target proteins. To avoid overexpression artifacts and ensure detection of proteins SUMOylated under physiologically relevant conditions, we immunopreciptated endogenous SUMO-conjugated proteins from HIV-1-infected versus uninfected HeLa cells using anti-SUMO1 and anti-SUMO2/3 antibodies. Both SUMO1- and SUMO2/3-conjugates were successfully enriched in the HIV-1-infected and uninfected samples (Figure 2B, C). In the SUMO1 IPs, SUMO1-RanGAP1 is by far the most abundant SUMO-protein conjugate and was used as a control given that its level of SUMOylation is unaffected by HIV-1 infection (Figure 2B). Anti-IgG IP was used as a negative control for all IPs.

**Figure 2.**
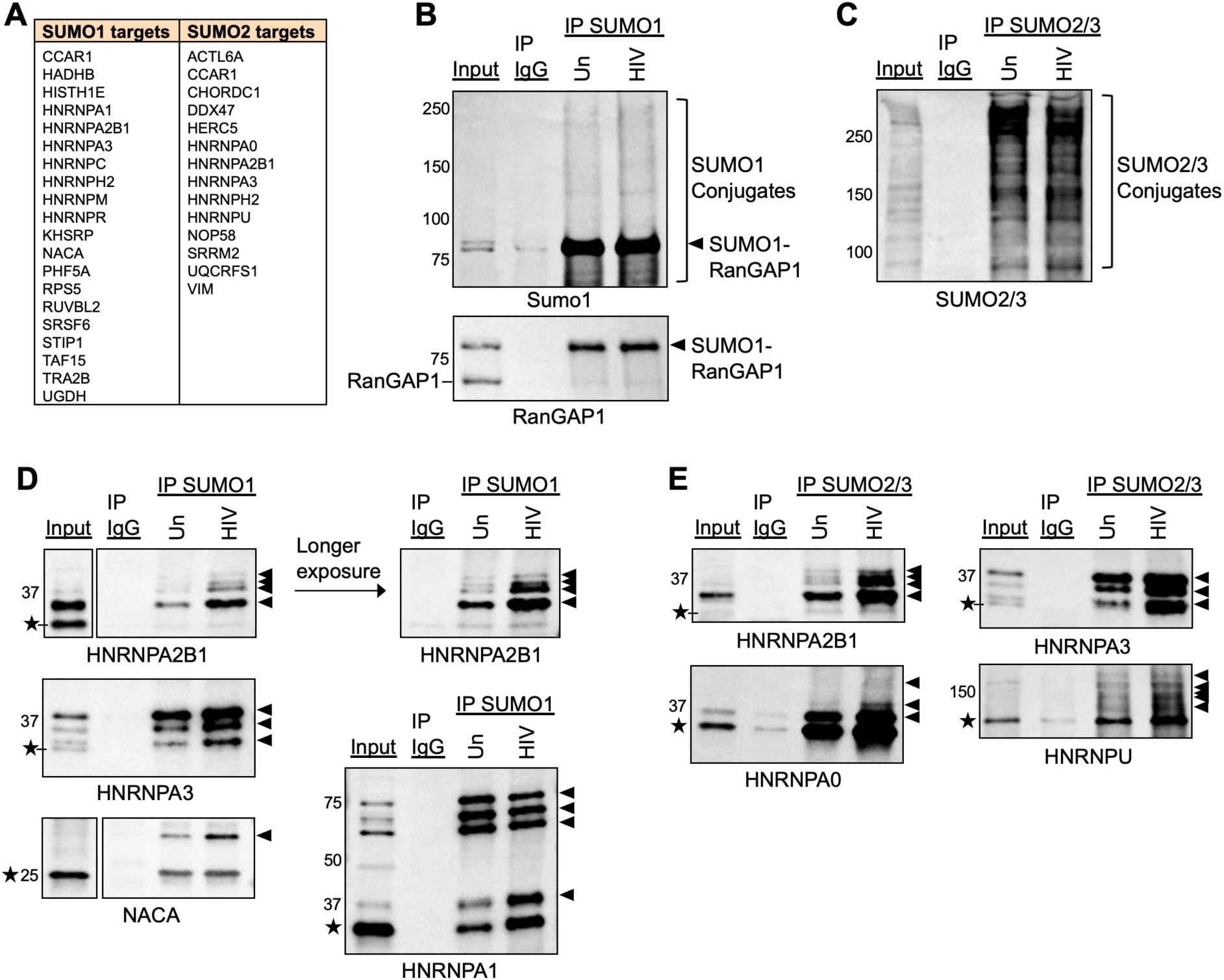
Experimental validation of HIV-1 induced SUMO targets. **(A)** List of putative HIV-1-induced SUMO1 and SUMO2 targets tested experimentally. **(B,C)** Western blots showing enrichment of SUMO1-conjugates, SUMO1-modified RanGAP1, and SUMO2/3-conjugates following immunoprecipitation with the indicated anti-SUMO antibodies in HIV-1-infected (“HIV”) versus uninfected controls (“Un”) at 24 hpi. **(D, E)** Western blot analysis of SUMO1- or SUMO2/3-enriched samples from HIV-1-infected cells (”HIV”) versus uninfected controls (“Un”). SUMO-conjugates were immunoprecipitated using anti-SUMO1 or SUMO2/3 antibodies at 24 hpi, and samples were analyzed by western blot for the indicated substrates. Stars indicate unmodified proteins. Arrowheads indicate SUMO-conjugated proteins. Crude lysate from HIV-1-infected cells was used as input for IPs, where input was 5% protein loaded in IPs. Non-immune IgG = IP control.

SUMO1- and SUMO2/3-enriched samples were then analyzed by western blot with antibodies against the endogenous candidate proteins of interest. Using this approach, we detected SUMO-modified forms of 12 SUMO1 targets and 13 SUMO2 targets, as evidenced by the presence of higher molecular weight SUMO-protein conjugates running above the unmodified proteins (Figure 2D, E, arrows, Supplemental Figure 3). Of these proteins, 4 SUMO1 targets (HNRNPA2B1, HNRNPA3, HNRNPA1, and NACA) and 4 SUMO2/3 targets (HNRNPA2B1, HNRNPA3, HNRNPA0, and HNRNPU) showed a clear increase in one or more SUMO-modified forms following HIV-1 infection, recapitulating our mass spectrometry data (Figure 2D, E). This HIV-1-induced increase in SUMOylation was particularly dramatic for HNRNPs A2B1 and A3 (Figure 2D, E). For both HNRNPs, infection with HIV-1 did not alter the expression level of the unmodified form of the protein (Supplemental Figure 2B), suggesting a specific increase in SUMOylation. Four of the SUMOylated HNRNPs, including A2B1, A3, A1, and A0, comprise the HNRNPA/B protein family, a structurally and functionally conserved subgroup involved in the regulation of transcription, alternative splicing, and translation.^33^ Thus, the observed SUMOylation of this protein family may represent a concerted approach used by the virus to manipulate essential stages of RNA metabolism upon infection.

A slight increase in SUMOylation following HIV-1 infection was also detected for 9 additional SUMO targets including HNRNPs M, C, and H2, as well as STIP1, DDX47, NOP58, CHORDC1, RPS5, and VIM (Supplemental Figure 3A). However, analysis for these proteins was complicated by low SUMOylation levels and the presence of nonspecific bands detected by the antibodies. Proteins that were SUMOylated basally regardless of HIV-1 infection status are shown in Supplemental Figure 3B. We were unable to detect SUMOylation of 9 proteins, suggesting that these proteins may represent false positives in our mass spectrometry data. Alternatively, given that only a small, 5-10% fraction of many proteins is SUMOylated at a given time,^34^ endogenous SUMOylation of these targets may be below our detection level.

Together, these data demonstrate that infection with HIV-1 leads to the SUMO-modification of multiple HNRNPs in both mass spectrometric and immunoprecipitation assays. We selected HNRNPA2B1 (A2/B1) and HNRNPA3 for further study given that both proteins are SUMOylated at a high level, and demonstrate a dramatic increase in their modification by both SUMO1 and SUMO2 paralogs upon HIV-1 infection. For the remainder of this text, we refer to these proteins as “A2B1” and “A3”.

### A2B1 and A3 are SUMOylated during HIV-1 infection

In our IP assays, HIV-1 infection led to an increase in four putatively SUMO1- or SUMO2/3- modified forms of A2B1: an abundant SUMO-modified form visible above the unmodified protein even in crude lysates (Figure 2D, E, Supplemental Figure 2B, arrows), and three less abundant, higher molecular weight SUMO-modified forms that were only detectable following enrichment of SUMO (Figure 2D, E, arrows). Here, the abundant form is likely monoSUMOylated A2B1, while the higher molecular weight bands may represent polySUMOylated forms of the protein. Unlike SUMO1, SUMO2/3 contains the SUMOylation consensus sequence and therefore forms chains readily. However, mixed chains with a terminal SUMO1 have been reported.^34,35,36^ Therefore the higher molecular weight forms of A2B1 observed in our assays may represent SUMO1-capped mixed chains. In our IP assays for A3, three SUMO-modified forms of A3 were detected above the unmodified protein even in crude lysates, and were selectively enriched following IP of SUMO1 or SUMO2/3 (Figure 2D, E, arrows). A clear increase in two of these forms was observed following HIV-1-infection, suggesting that although A3 is highly SUMOylated basally, infection with HIV-1 leads to an increase in two specific SUMOylated forms of the protein (Figure 2D, E, arrows).

To confirm that the novel species were indeed due to SUMOylation, we examined the effects of the SUMOylation inhibitor TAK-981^37^ on the SUMOylation of A2B1 and A3 during HIV-1 infection. TAK-981 treatment was confirmed to dramatically reduce the SUMOylation of RanGAP1 relative to the DMSO control, indicating successful inhibition of SUMOylation (Figure 3A). SUMO1-conjugates were immunoprecipitated from HIV-1-infected cells treated with or without TAK-981, followed by western blot analysis with anti-A2B1 and A3 antibodies. SUMO1 IP samples from uninfected, DMSO-treated cells were used as a control. Here, TAK-981 treatment prevented the HIV-1-induced SUMOylation of both A2B1 and A3, as observed by reduced levels of all SUMOylated forms discussed above relative to the HIV-1-infected, DMSO control (Figure 3B). The level of unmodified A2B1 and A3 was not affected by TAK-981 treatment (Figure 3B). Together, these results suggest that the bands observed in our IP assays represent bonafide SUMO-modified forms of A2B1 and A3. Additionally, IP analysis in the opposite direction (IP of A2B1, followed by western blotting with anti-SUMO1 and anti-SUMO2/3 antibodies) confirmed HIV-1-induced SUMOylation of A2B1 (Supplemental Figure 4A).

**Figure 3.**
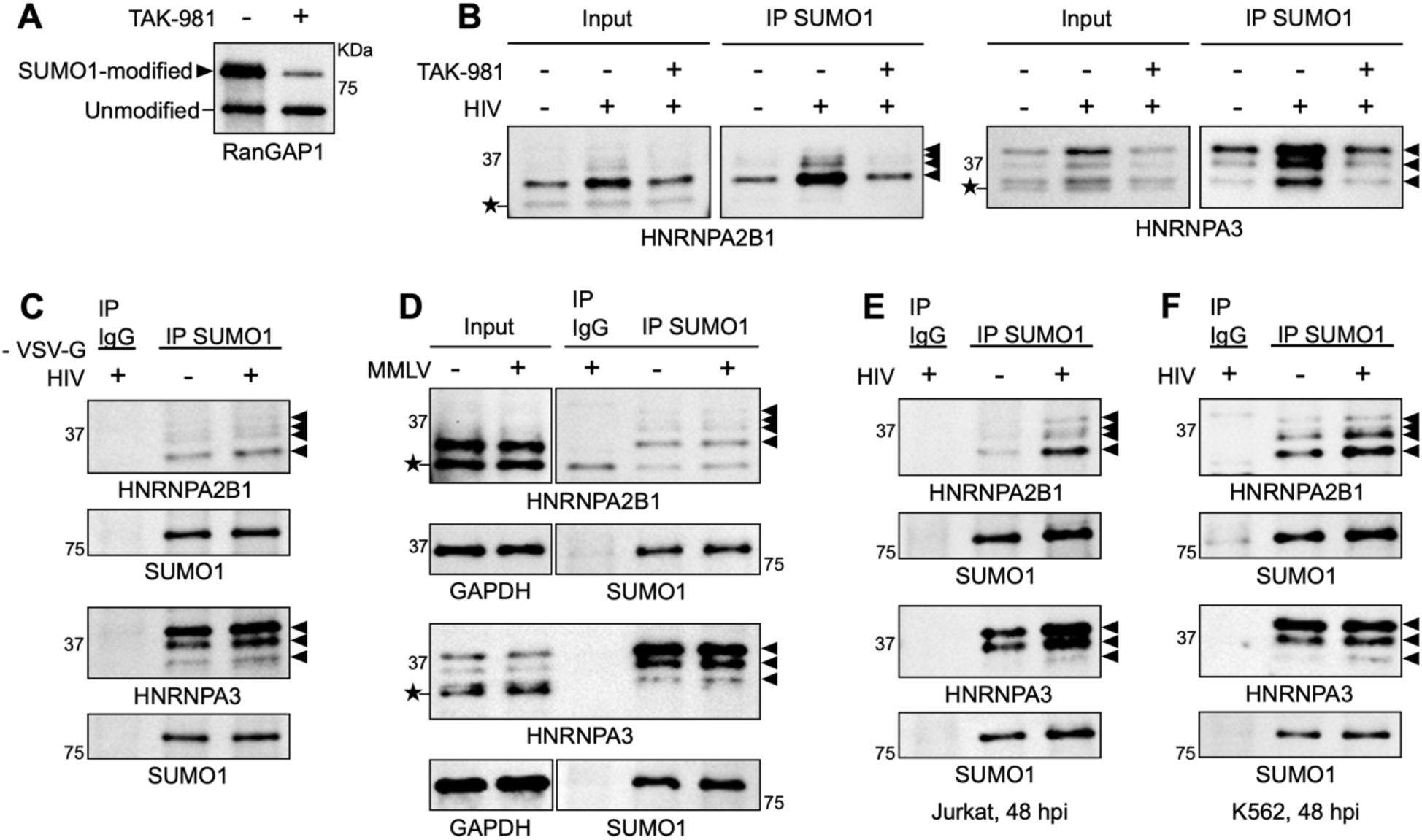
HNRNPs A2B1 and A3 are SUMOylated during HIV-1 infection. **(A)** Western blot showing the level of SUMO1-modified and unmodified RanGAP1 in HeLa cells treated with the SUMOylation inhibitor TAK-981 versus the DMSO control (“-”). **(B)** Western blot showing the level of SUMO1-conjugated A2B1 and A3 in HIV-1-infected cells treated with or without TAK-981. Uninfected, untreated cells were used as a control. Crude lysate from the corresponding conditions was used as input for IPs (5% of total protein loaded in IPs). For A and B, TAK-981 concentration = 12 µM, DMSO = untreated control. **(C)** Western blot analysis of SUMO1-modified A2B1 and A3 in cells infected with Env mutant HIV-1 (pNL4-3 deltaENV-EGFP) lacking the VSV-G envelope (“-VSV-G HIV”) versus the uninfected control. Cells were harvested at 48 hpi. **(D)** Western blot analysis of SUMO1-modified A2B1 and A3 in MMLV-infected HeLa cells versus uninfected controls. Cells were infected with VSV-G pseudotyped MMLV (pNCA-GFP) and harvested at 48 hpi for lysate preparation. Crude lysate input for each condition is shown for comparison. **(E, F)** Western blot analysis of SUMO1-modified A2B1 and A3 in HIV-1-infected Jurkat cells (E) or K562 cells (F) versus uninfected controls at 48 hpi. For all, SUMO1-conjugated proteins were immunoprecipitated using anti-SUMO1 antibodies and samples were analyzed by western blot for A2B1 and A3. Arrows indicate SUMO1-modified proteins. Stars indicate unmodified proteins. For C-F, non-immune IgG = IP control and SUMO1-modified RanGAP1 = loading control for IPs.

### HIV-1 drives the SUMOylation of A2B1 and A3 in multiple cell line models

We next sought to characterize the requirement of the observed SUMOylation phenotype for HIV-1. We first infected cells with an HIV-1-GFP reporter virus lacking a functional envelope protein (Figure 3C, “-VSV-G HIV”). Infection with this virus did not lead to increased A2B1 and A3 SUMOylation, indicating that the SUMO-modification of these proteins is dependent on successful entry of the HIV-1 virion into the host cell (Figure 3C). Additionally, SUMOylation of A2B1 and A3 was not observed upon infection with the retrovirus Moloney Murine Leukemia Virus (MMLV), despite comparable infection efficiencies (Figure 3D, Supplemental Figure 4B), suggesting that the SUMOylation of A2B1 and A3 is induced specifically by HIV-1.

To explore the range of cells responding to HIV-1 infection with increased A2B1 and A3 SUMOylation, we repeated our immunoprecipitation assays in additional cell lines. HIV-1-driven SUMOylation of A2B1 and A3 was also observed in Jurkat cells (an immortalized human T lymphocyte line) and in K562 cells (a chronic myelogenous leukemia cell line), though this phenotype was less dramatic in the K562 cells (Figure 3E, F).

### SUMOylation of A2B1 and A3 requires HIV-1 integration

We next examined the time course of A2B1 and A3 SUMOylation during viral infection. SUMO1-conjugates were immunoprecipitated from HIV-1-infected and uninfected HeLa cells at 6, 12, 24, and 48 hours post infection (hpi), followed by western blot analysis with anti-A2B1 and A3 antibodies (Figure 4A, B). At 6 and 12 hpi, there was no difference in the level of SUMOylated A2B1 between the HIV-1-infected samples versus the uninfected controls. However, at both 24 and 48 hpi, increased SUMOylation of A2B1 was readily apparent in the HIV-1-infected samples (Figure 4A). Similarly, the highest level of SUMOylated A3 was detected in the HIV-1-infected samples at 48 hpi, although there was a slight increase in the level of one SUMO1-A3 conjugate as early as 6 hpi (Figure 4B). Integration of the HIV-1 DNA into the host cell genome reportedly occurs at ∼15 hpi.^38^ Thus, the observed timing of HIV-1-induced SUMOylation of A2B1 and A3 suggests that this modification occurs post-integration. Consistently, infection with integrase-competent HIV-1 virus was required to induce maximum SUMO-modification of A3 and A2B1, in contrast to the respective integrase-deficient condition^39,40^ (Figure 4C). Together, these findings suggest that SUMOylation of A2B1 and A3 depends upon viral gene products expressed after integration.

**Figure 4.**
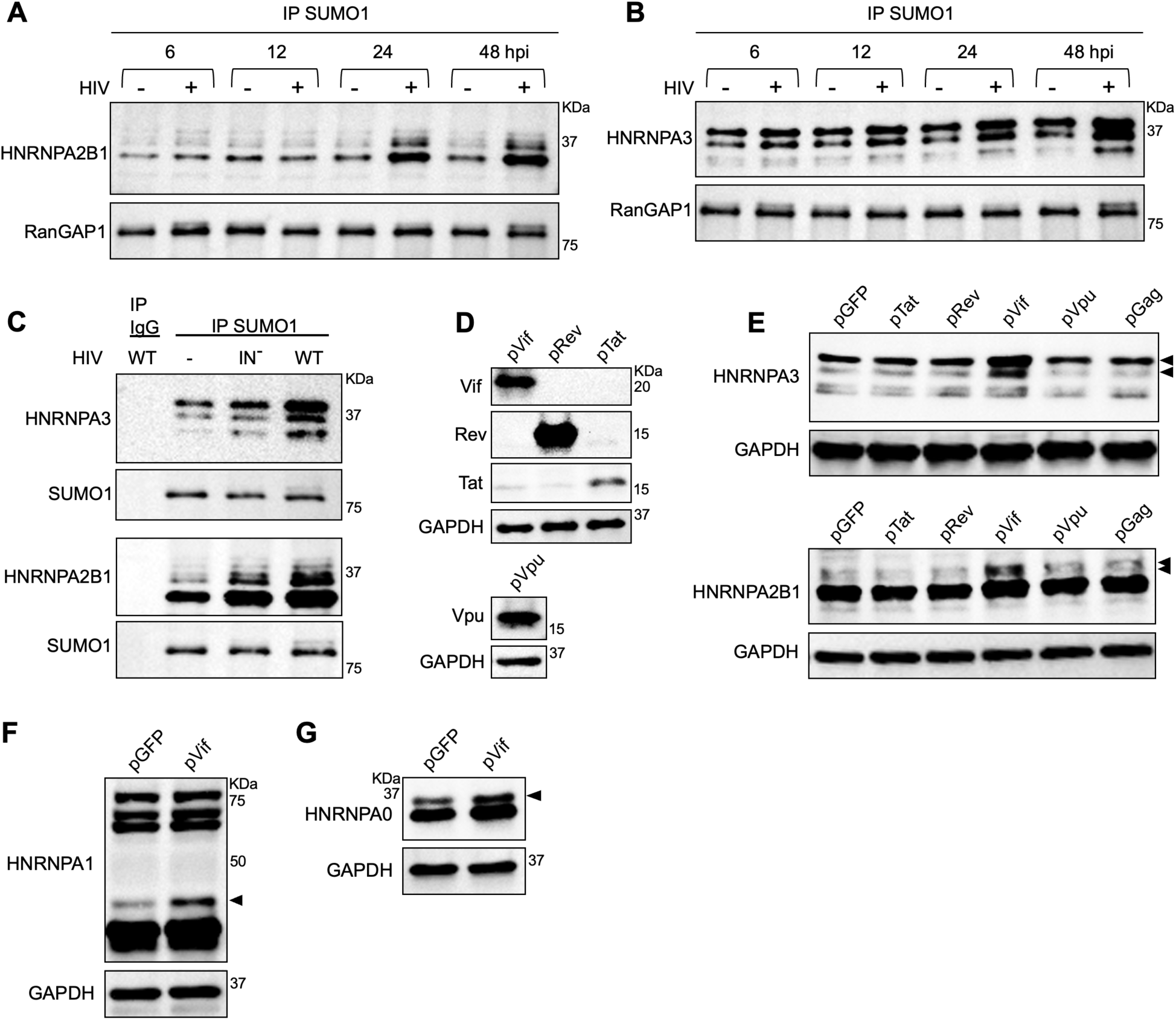
Expression of HIV-1 viral protein Vif drives the SUMOylation of HNRNPs A2B1 and A3. **(A,B)** Western blots showing the level of SUMO-modified A2B1 or A3 in HIV-1-infected cells versus uninfected controls at 6, 12, 24, and 48 hpi. **(C)** Western blot analysis of SUMO1-modified A3 and A2B1 in cells infected with integrase-deficient HIV-1 (IN^-^) versus the respective integrase WT condition (pNL4-3.ZsGreen.R−.E− IN-D64A versus pNL4-3.ZsGreen.R−.E− IN-wt, respectively).^39,40^ “-” = uninfected control. Non-immune IgG = IP control. For A-C, SUMO1-conjugated proteins were immunoprecipitated using anti-SUMO1 antibodies and samples were analyzed by western blot for A2B1 or A3. SUMO1-modified RanGAP1 = loading control for IPs. **(D)** Western blot showing expression of the HIV-1 viral proteins Vif, Rev, Tat, and Vpu in cells transfected with a plasmid expressing each protein. Plasmids used are detailed in Methods and Materials. **(E)** Western blot analysis of SUMO-modified A3 and A2B1 (arrows) detected following transfection of pTat, pRev, pVif, pVpu, and pGag. **(F,G)** Western blot analysis of SUMO-modified HNRNPA1 (F) or HNRNPA0 (G) following transfection of pVif. Arrows indicate SUMO-modified forms also shown to increase upon HIV-1 infection in Figure 2. For parts E-G, a plasmid expressing only GFP (pGFP) = transfection control. GAPDH = loading control. For C-E, cells were harvested at 48 hpi (C), or 48 hours post-transfection (D-G). HeLa cells were used for A-G.

### Vif drives the SUMOylation of A2B1 and A3

We next tested whether expression of a specific HIV-1 viral protein induces the SUMOylation of A2B1 and/or A3. The HIV-1 viral genome encodes structural proteins Gag, Gag-Pro-Pol, and Env, regulatory proteins Tat and Rev, and accessory proteins Nef, Vif, Vpr, and Vpu. In principle, any of these gene products could induce A2B1 and A3 SUMOylation. Infection with pseudotyped virions delivering HIV-1 genomes unable to express Env (Figures 1 and 2, pNL4-3 deltaENV-EGFP), or Env, Vpr, and Nef (Figure 4C, pNL4-3.ZsGreen.R−.E−), still resulted in induced SUMOylation of A2B1 and A3, suggesting that these proteins are dispensable for the observed SUMOylation. Thus, we focused our attention on the remaining proteins Gag, Vif, Vpu, Rev, and Tat. Cells were transfected with plasmids expressing each of these proteins and expression of the respective protein was verified by western blot (Figure 4D). The induction of SUMOylated A2B1 and A3 were monitored by the presence of higher molecular weight SUMO-conjugates running above the unmodified proteins on western blot. Only expression of Vif led to the appearance of multiple SUMOylated forms of A3 and A2B1 (Figure 4E, arrows), whereas expression of Tat, Rev, Vpu, and Gag showed no induction of A3 and A2B1 SUMOylation relative to the control. These findings indicate that the HIV-1 viral protein Vif drives the SUMOylation of A2B1 and A3 during infection, suggesting a novel function for this protein aside from its well-studied role in mediating the ubiquitination of APOBEC3G.^9^ Vif was able to induce the SUMOylation in the absence of any other viral proteins.

### Vif drives the SUMOylation of the HNRNPA/B family

HNRNPA1 and HNRNPA0, the remaining members of the structurally-conserved HNRNPA/B family^41^, were also found to be SUMOylated upon HIV-1 infection (Figure 1E, F, Figure 2D, E). We next asked whether Vif also drives their SUMOylation. Indeed, expression of Vif led to an increase in the specific SUMOylated form of HNRNPA1 that was also found to increase following HIV-1 infection (Figure 4F, arrow, as compared to Figure 2D). Similarly, Vif expression led to an increase in the most predominant SUMOylated form of HNRNPA0 (Figure 4G, arrow, as compared to Figure 2E). Together, our results suggest a common Vif-mediated mechanism driving the SUMOylation of the entire HNRNPA/B family. These findings raise the intriguing possibility that this modification may remodel HNRNPA/B family function upon infection, in turn mediating their role in HIV-1 RNA metabolism.

### Depletion of A2B1 and A3 alters HIV-1 splicing

Alternative splicing is critical for the production of infectious HIV-1 virions: splicing of the HIV-1 primary transcript greatly expands the coding capacity of its compact viral genome, allows temporal regulation of HIV-1 gene expression, and is necessary for the expression of most HIV-1 viral proteins.^42,43,44,45^ In infected cells, unspliced (US) viral transcripts encode Gag and Gag-Pro-Pol proteins and serve as the genomic RNA packaged into virions. HIV-1 RNAs can also undergo alternative splicing, where spliced transcripts include a 4-kb class of incompletely spliced (IS) RNAs, and a 2-kb class of completely spliced (CS) RNAs^42,43,44,45^ The pattern of HIV-1 splicing has been extensively described and occurs through the use of four splice donor sites (D1-4) and ten splice acceptors (A1-3, A4a, A4b, A4c, A4d, A5, A5b, A7) distributed along the HIV-1 genome. IS RNAs result from splicing between the D1 donor site and one of several splice acceptors: A1 for transcripts encoding Vif, A2 for Vpr, and A4 or A5 for Env/Vpu. CS RNAs result following a second splicing event between D4 and A7, and encode Tat, Rev, or Nef.^42,43,44,46,47,48^ HIV-1 splice site usage is controlled by both *cis*-acting splicing regulatory elements (SREs) and *trans*-acting cellular RBPs, including HNRNPA/B family members.^47,49,50^

To examine the role of HNRNPs A2B1 and A3 in mediating proper HIV-1 RNA splicing, we utilized a previously described Primer ID-tagged deep sequencing assay^46^ to quantify the effects of A2B1 and A3 depletion on the ratio of US/IS/CS HIV-1 RNAs, and HIV-1 splice acceptor site usage. We knocked down expression of A2B1 and A3 in HeLa cells (Figure 5A) followed by infection with VSV-G-pseudotyped HIV-1 (pNL4-3 deltaENV-EGFP). Loss of HNRNPA2B1 and HNRNPA3 had no significant effect on the ratio of US versus total spliced RNAs (Figure 5B). However, within the spliced classes (comprised of both IS and CS RNAs), loss of HNRNPA3 led to a dramatic increase in IS RNAs over CS RNAs (Figure 5B’). Additionally, loss of HNRNPA2B1 led to significantly increased A2 and A3 splice site usage, and a decrease in the use of the A1 splice site (Figure 5C, 5D). In contrast, loss of HNRNPA3 led to decreased use of the A2 and A4 splice sites (Figure 5C, 5D). These results suggest that both HNRNPs play significant roles in regulating the relative proportions of the various HIV-1 mRNAs late in the viral life cycle.

**Figure 5.**
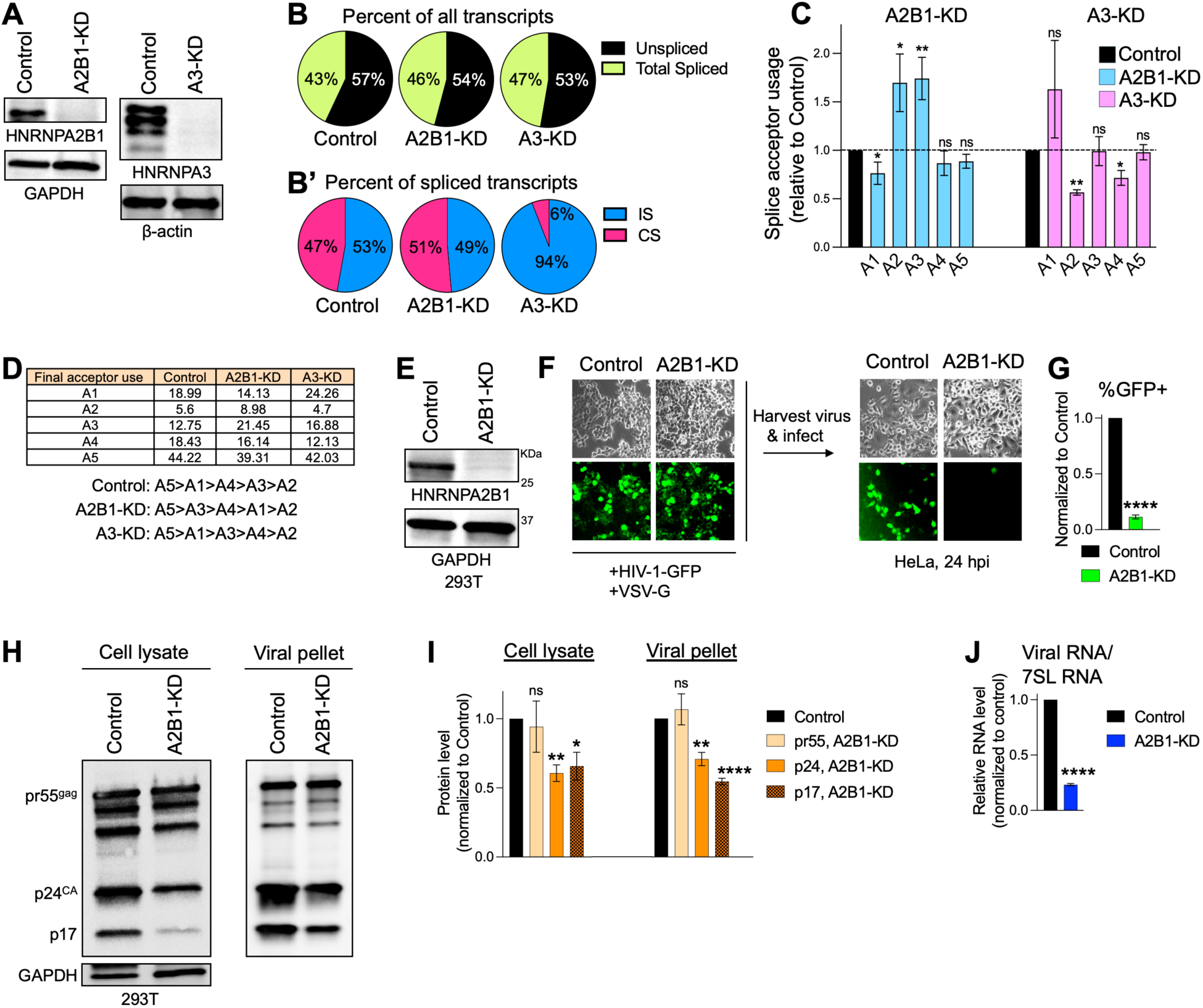
Depletion of HNRNPs A2B1 and A3 alters HIV-1 splicing and reduces infectivity. **(A)** Western blot showing A2B1 and A3 expression in HeLa cells transfected with Control, A2B1, or A3 siRNAs. **(B)** Quantification of Unspliced and total Spliced HIV-1 RNAs in siRNA Control versus A2B1- or A3-knockdown (KD) cells. Circle proportions represent percent of all transcripts. Cells were infected with VSV-G pseudotyped HIV-1-GFP reporter virus and splicing was quantified at 24 hpi. **(B’)** Quantification of incompletely spliced (IS) and completely spliced (CS) HIV-1 RNAs in Control vs. A2B1-KD and A3-KD samples at 24 hpi. Circle proportions represent percent of spliced transcripts. **(C)** Analysis of final splicing events occurring at HIV-1 splice acceptors A1 through A5 in A2B1-KD and A3-KD samples. Splice acceptor usage is represented as fold change relative to the siRNA Control. **(D)** Table showing average percent of spliced transcripts utilizing A1 through A5 final splice acceptors in Control vs. A2B1- and A3-KD samples. The order of splice acceptor use, from most to least common, is listed for each sample. **(E)** Western blot showing A2B1 expression in Lenti-X 293T cells transfected with Control versus A2B1 siRNAs. **(F)** Control and A2B1-KD cells from (E) were co-transfected with plasmids expressing the HIV-1-GFP reporter genome and the VSV-G envelope. Cell culture supernatants were collected at 48 hours post-transfection and used to infect HeLa cells. **(G)** Flow cytometric analysis of the percent GFP+ HeLa cells from (F) at 24 hpi. **(H)** Western blot showing expression of HIV-1 Gag cleavage products in Control versus A2B1-KD 293T cells (Cell Lysate) and in released virions (Viral Pellet) after transfection with plasmids expressing the HIV-1 viral genome and VSV-G envelope. Cells were harvested for lysate preparation at 48 hpi and virions in the supernatants were isolated by pelleting through a 25% sucrose cushion. **(I)** Densitometric analysis of data from (H) shown relative to Control. **(J)** RT-qPCR analysis of HIV-1 viral RNA in the virions released from Control versus A2B1-KD cells. 7SL RNA was used as an internal control. (error bars = mean ± SEM of three independent experiments; for splicing analysis, siRNA Control and A2B1-KD sample data was derived from two independent experiments each with three technical replicates, data for A3-KD samples includes two technical replicates from one experiment; unpaired t-test,****p≤0.0001,***p≤0.001,**p≤0.01,*p≤0.05).

### Loss of A2B1 inhibits production of infectious HIV-1 virus

We next examined the role of A2B1 and A3 in the production of infectious HIV-1 particles. Control and A2B1-KD Lenti-X 293T cells (Figure 5E) were transfected with plasmids expressing the HIV-1-GFP reporter virus and the vesicular stomatitis virus (VSV-G) envelope glycoprotein (Figure 5F). Viral supernatants were harvested at 48 hours post-transfection and equal volumes of undiluted supernatants were used to infect naïve HeLa cells (Figure 5F). Virus produced from A2B1-KD cells displayed dramatically reduced infectivity (%GFP+) in comparison to control virus (Figure 5F, G, Supplemental Figure 5A). Unlike A2B1-KD, KD of A3 led to substantial loss of 293T cell viability (data not shown). Thus, the effect of A3-KD on virus production was not assessed.

To test whether the reduced infectivity of virus produced by the A2B1-KD cells was accompanied by alterations in HIV-1 protein expression, we examined the levels of Gag protein in the cells (cell lysate) versus their released virions (viral pellet) (Figure 5H, I). Intriguingly, A2B1-KD had no effect on the level of Pr55^gag^ in the virus-producing cells or virions. However, the Pr55^gag^cleavage products p24^CA^ and p17 were reduced in both the A2B1-KD cells and released virions relative to the control (Figure 5H, I), suggesting that A2B1-KD may alter the processing of Gag.

During HIV-1 virus production, US HIV-1 RNAs are packaged into assembling virions. To test the effect of A2B1 loss on HIV-1 RNA packaging, we extracted total RNA from pelleted virions released from control and A2B1-KD 293T cells, and compared the level of HIV-1 RNA by RT-qPCR (Figure 5J). Cellular 7SL RNA is abundantly packaged into HIV-1 virions and was used as an internal control.^51^ We observed a dramatic reduction in HIV-1 RNA in the A2B1-KD virions relative to the control (Figure 5J), whereas the level of 7SL RNA in the viral pellet was unaffected by A2B1 KD.

All together, these findings suggest that the effect of A2B1 depletion on virus production is likely multifactorial and occurs through a combination of effects on alternative splicing, export, translation, and/or the packaging of HIV-1 RNAs into virions. This is consistent with the myriad roles of A2B1 in regulating multiple aspects of RNA metabolism, and raises the possibility that the HIV-driven SUMOylation of A2B1 may modulate one or more of these regulatory roles.^32,33^

## DISCUSSION

As intracellular parasites, HIV-1 and other viruses are entirely dependent upon the molecular pathways of the host cell for completion of their viral life cycles. Viral exploitation of post-translational modifications, including SUMOylation, provides an opportunity to fine tune these host pathways, “rewiring” them to the viruses’ advantage.^2,3,6,8,9,10,11,52,53^ In this study, we utilized quantitative proteomics to characterize the effect of HIV-1 infection on the SUMO-modification status of the host cellular proteome. This work represents the first large-scale, unbiased proteomic screen for SUMOylated substrate proteins in the setting of HIV-1 infection. We present evidence that infection with HIV-1 leads to a reprogramming of host cell SUMOylation, resulting in a striking pattern of widespread increased SUMO-modification of proteins involved in alternative splicing, and RNA metabolism broadly (Figure 1E-H). These proteins included numerous heterogeneous nuclear ribonucleoproteins (HNRNPs), and all members of the structurally conserved HNRNP A/B protein family (A2B1, A3, A1, and A0), for which HIV-driven SUMOylation was experimentally validated in multiple biochemical assays with the endogenously-expressed proteins (Figure 2D, E, Figure 3A-F).

An intriguing aspect of the HNRNPA/B family’s SUMOylation is that it was found to be induced by the post-integration expression of the HIV-1 Viral Infectivity Factor (Vif) (Figure 4C-E), where expression of Vif alone led to an increase in the specific SUMOylated forms of these proteins that were also increased following infection with the HIV-1 virus (Figure 4E-G). Vif has an extensively characterized role in mediating the poly-ubiquitination and subsequent proteasomal degradation of APOBEC3G, a host restriction factor that impairs HIV-1 replication by triggering hypermutation of the viral reverse transcripts.^54,55,56,57,58^ Ubiquitination of APOBEC3G is achieved through the formation of a Cullin-RING E3 ubiquitin ligase complex (CRL5) composed of Cullin5 (Cul5), ElonginB (EloB), ElonginC (EloC), and RING-box protein 2 (RBX2) in the presence of Vif and the host co-factor CBF-β.^9^ This E3 ubiquitin ligase complex and its components are not known to have concomitant E3 SUMO ligase activity. Thus, our findings of Vif-driven HNRNPA/B SUMOylation suggest a novel function for Vif, perhaps mediated by its interaction with a yet-to-be identified E3 SUMO ligase. Interestingly, the interaction of Vif with APOBEC3G has been shown to be RNA-mediated,^59,60^ suggesting that a Vif-E3 SUMO ligase complex may associate with HIV-1 RNA bound to the HNRNPA/B proteins. Alternatively, HNRNPA/B SUMOylation may be indirect and dependent on the degradation of APOBEC3G. During infection with human adenovirus type 5 (HAdV-C5) for instance, APOBEC3A decreases the SUMOylation of the adenoviral DNA-binding protein E2A,^61^ raising the possibility of an indirect mechanism where Vif-mediated degradation of APOBEC3G releases its inhibition of HNRNPA/B SUMOylation.

A central question that remains is the functional consequence of the HIV-induced, Vif-driven SUMOylation. Multiple HNRNPs, including but not limited to the HNRNPA/B family, regulate the alternative splicing of HIV-1 viral pre-mRNAs, a process critical for viral replication.^47,50^ The relative usage of HIV-1 splice sites is in part regulated by the binding of HNRNPs to *cis*-acting splicing regulatory elements located within the HIV-1 RNAs. In particular, HNRNPA/B proteins have been shown to module the splicing of HIV-1 RNAs by binding to exonic splicing silencers (ESSs) located within Tat exon 2, Tat/Rev exon 3, the Vif exon, or an intronic splicing silencer (ISS) located within Tat intron 2.^50,62,63,64,65,66^ Consistently, we have shown that genetic depletion of HNRNPs A2B1 and A3 led to distinct patterns of altered HIV-1 splice acceptor site usage (Figure 5C,D) and/or altered ratios of US/IS/CS HIV-1 RNAs (Figure 5B, B’). These findings were accompanied by dramatically reduced the infectivity of the virions released by the A2B1-depleted cells (Figure 5F, G).

Moreover, the predicted SUMOylated K residues of the HNRNPA/B proteins are overwhelmingly concentrated within their RNA-binding domains. HNRNPA/B proteins contain two N-terminal RNA recognition motifs (RRM1 and RRM2) known to mediate RNA-binding, and a C-terminal low complexity domain (LCD), known to primarily mediate protein-protein interactions (Supplemental Figure 6A, B).^41^ We utilized SUMOplot^TM^ to predict the SUMOylated sites within A2B1 and A3 (Supplemental Figure 6A’, B’). SUMOplot^TM^ scores the probability of SUMOylation at a given K based on similarity to the SUMO consensus motif ΨKxD/E (where Ψ is a hydrophobic residue and x is any amino acid), and known SUMOylated sites in other proteins.^67^ Intriguingly, ∼70% of the predicted SUMO sites are located within the RRMs of each protein (Supplemental Figure 6A-C). These predicted SUMO sites include K108, the residue of A2B1 that has been shown to be SUMOylated in multiple prior studies.^41,68,69^ Similarly, there is dramatic enrichment of predicted SUMO sites in the RRMs of the remaining HNRNPA/B family members, HNRNPA0 and HNRNPA1 (Figure 6C).

**Figure 6.**
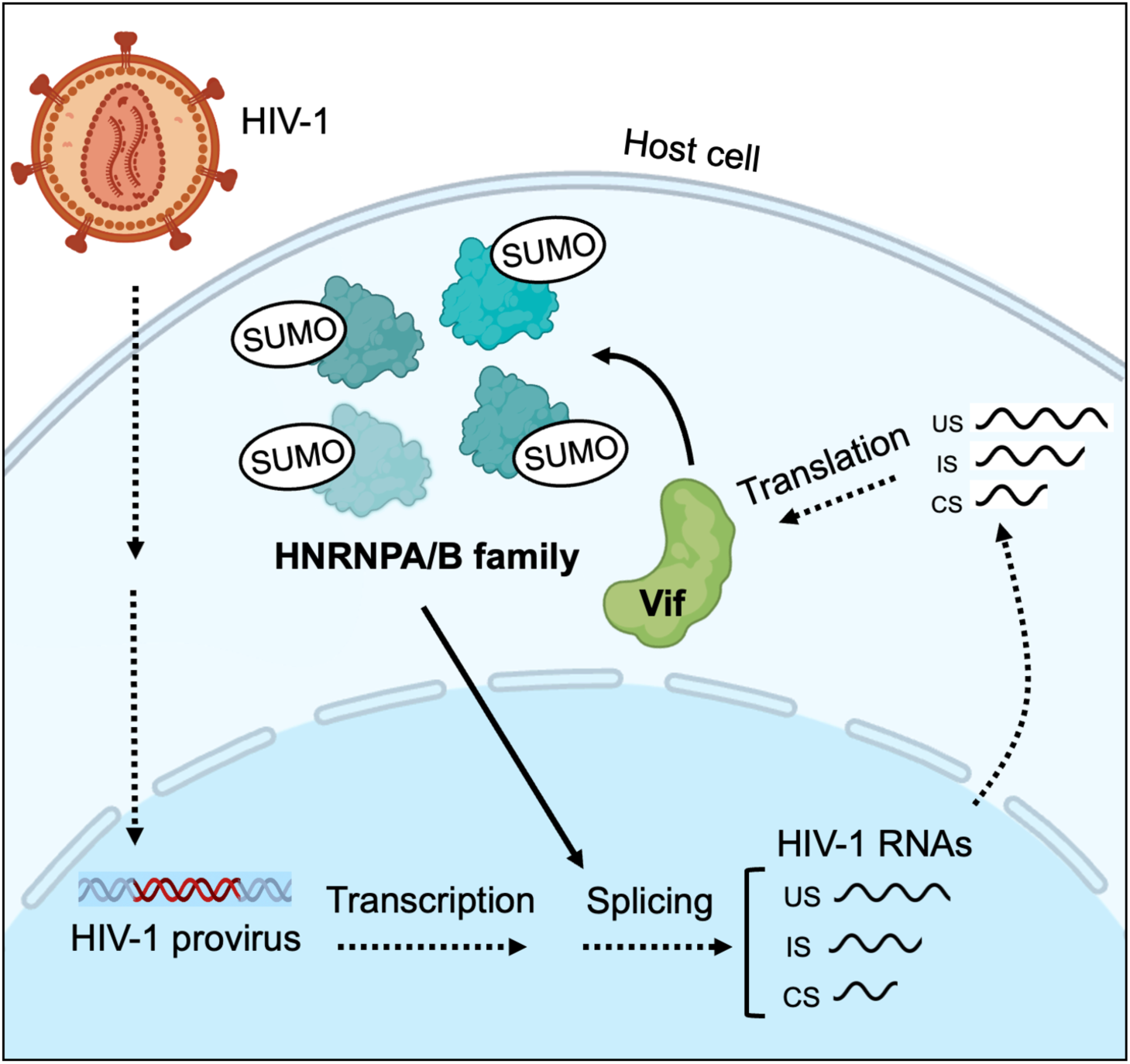
Graphical abstract. Expression of the HIV-1 viral protein Vif from the integrated provirus drives SUMOylation of the HNRNPA/B protein family, likely modulating the alternative splicing of HIV-1 viral mRNAs.

The RRMs of HNRNPA/B proteins cooperatively mediate interactions with specific RNA sequence motifs and are critical for their function as regulators of alternative splicing.^12,41^ PTMs regulate the activities of numerous HNRNPs, where SUMOylation may affect splicing activity by altering RNA-binding by direct steric hindrance, or interaction with other splicing factors and spliceosome components.^11,70,71^ All together, these findings suggest that HIV-driven SUMOylation of the HNRNPA/B family may alter their binding to regulatory elements within the HIV-1 mRNAs, in turn modulating HIV-1 RNA splicing. Given that the HNRNPA/B family is canonically central to the regulation of mammalian alternative splicing,^33,41^ our findings also raise the intriguing possibility that HIV-driven HNRNPA/B SUMOylation may simultaneously remodel the splicing of host transcripts in HIV-infected cells.

Of important note, HNRNPA/B proteins also regulate stages of RNA metabolism aside from alternative splicing, including the transcription, transport, stability, and translation of mRNAs.^33^ In HIV-1 infected cells, A2B1 and A1 influence HIV-1 RNA export.^72,73,74^ SUMOylation of A2B1 has also been shown to mediate the sorting of miRNAs into exosomes^75^ and the loading circRNAs into extracellular vesicles,^69^ an intriguing finding given the impaired HIV-1 viral RNA packaging we observed following loss of A2B1 (Figure 5J). Therefore, HIV-driven HNRNPA/B SUMOylation may simultaneously modulate their role in multiple aspects of viral mRNA production. Future mutagenesis analysis and the generation of un-SUMOylatable HNRNPA/B mutants will provide powerful tools to dissect the precise consequences of this modification, both for HIV-1 and host cellular RNA metabolism.

In summary, our study has uncovered a novel mechanism by which infection with HIV-1, and the subsequent expression of the HIV-1 viral protein Vif, induces the concerted SUMOylation of the HNRNPA/B family of RNA-binding proteins. These findings suggest a previously unappreciated function for Vif, and have significant implications for the regulation of viral and host cellular alternative splicing in HIV-infected cells.

## METHODS AND MATERIALS

### Cell culture and cell lines

HeLa and HEK293T cells were maintained in DMEM (Gibco #11965092) supplemented with 10% fetal bovine serum (FBS), 100 units/mL Penicillin-Streptomycin (Gibco #15140122), and 2 mM L-Glutamine (Gibco #25030081). Jurkat cells were maintained in RPMI 1640 medium (Gibco #22400089) supplemented with 10% FBS. All cells were maintained at 37°C in a humidified incubator with 5% CO_2_. To generate cell lines stably expressing tagged C-terminal mutant SUMO proteins, 3xFlag-SUMO1^Q92R^, 3xFlag-SUMO2^Q88R^, 6His-SUMO1^Q92R^, and 6His-SUMO2^Q88R^ expression vectors were transfected into HeLa cells using Lipofectamine 3000 (Invitrogen #L3000075). Pooled cells stably expressing the desired proteins were selected using 1 mg/mL Geneticin (Gibco #10131035) and validated by western blot (WB).

### Virus production

Lenti-X 293T cells were seeded at 1×10^6^ cells per 10 cm dish one day prior to transfection. Cells were transfected with 15 µg of the pNL4-3 deltaENV-EGFP reporter (Centre for AIDS reagents #100616) and 5 µg of pCMV-VSV-G (Addgene #8454) using Lipofectamine 3000 (Invitrogen #L3000075) in serum-free DMEM. 6 hours later, transfection media was replaced with serum-containing DMEM. Viral supernatant was harvested at 48 hours post-transfection, filtered through a 0.45 µm filter, treated with 35 units/mL DNase (Invitrogen #AM2238) at 37°C for 45 min. Viral supernatant was either used directly for infection as described below, or virions in the supernatant were pelleted through a 25% sucrose cushion.

### Infection

3×10^6^ HeLa cells were seeded in 15 cm plates the day prior to infection. Jurkat cells were seeded the day of infection at 5×10^6^ cells per well of a 6 well plate. HeLa and Jurkat cells were incubated with 10 mL or 2 mL viral supernatant, respectively, plus 8 µg/mL Polybrene (Sigma #TR1003G). Virus was removed after 5 hours and replaced with fresh medium. Jurkat cells were resuspended in 10 mL fresh medium, and moved to 10 cm plates until harvest.

### Sample preparation for MS

3xFlag-SUMO1^Q92R^, 3xFlag-SUMO2^Q88R^, 6His-SUMO1^Q92R^, and 6His-SUMO2^Q88R^ cell lines were seeded and infected with HIV-1 in 15 cm plates as described above. One 15 cm plate was used for each sample and three replicates of each condition were prepared, including uninfected controls. Cells were harvested at 24 hpi and pelleted by centrifugation (1100 rpm for 5 min.). For 3xFlag-tagged samples, cells were resuspended in 150 µL SDS lysis buffer pre-heated to 95°C (1% SDS, 50 mM Tris/HCl pH 8.0, 150 mM NaCl). Samples were heated at 95°C for 10 min. 500 µL of RIPA w/o SDS (50 mM Tris/HCl pH 8.0, 150 mM NaCl, 1.1% Triton X-100, 0.55% Sodium deoxycholate) was added to each sample, followed by sonication (30-50% power for 3 x 1 sec.) until samples were no longer viscous. 850 µL RIPA w/o SDS was added to each sample, debris were pelleted by centrifugation (13.2K rpm for 10 min. at 4°C), and supernatants transferred to fresh tubes. 40 µL of Anti-FLAG M2 Magnetic Beads (Sigma, #M8823) were pre-washed 3x with RIPA (50 mM Tris/HCl pH 8.0, 150 mM NaCl, 1% Triton X-100, 0.5% Sodium deoxycholate, 0.1% SDS) and then added to each sample. Lysates were incubated with beads overnight at 4°C with rotation. Beads were then washed 3x with RIPA, and then 3x with RIPA w/o detergent (50 mM Tris/Hcl pH 8.0, 150 mM NaCl). Bound proteins were eluted in 30 µL elution buffer containing 50 mM Tris/HCL pH 8.0, 150 mM NaCl, 0.05% RapiGest SF Surfactant (Waters) and 100 µg/mL 3xFLAG peptide (MDYKDHDGDYKDHDIDYKDDDDK, Elim Biopharm) for 30 min. at room temperature on a shaker. 6His-tagged samples were prepared as described.^4^ Enrichment of Flag and His-tagged proteins were confirmed by WB of input, eluates, and flowthrough. Eluates were used below for both total protein analysis and identification of SUMO-conjugated peptides.

### LC-MS/MS

Immunoprecipitated samples were diluted in buffer (2M urea, 10 mM NH_4_HCO_3_, and 2 mM DTT) and incubated at 60°C for 30 min. Iodoacetamide was added to 2 mM and samples were incubated at room temperature in the dark for 45 min. 160 ng trypsin (Gold Mass spectrometry Grade, Promega) was added and samples incubated overnight at 37 °C. Samples were desalted with C18 spin columns (Nest Group, product #HUMS18R), vacuum centrifuged to dryness, and resuspended in 0.1% formic acid for MS analysis. All samples were analyzed on an Orbitrap Eclipse MS system equipped with an Easy nLC 1200 ultra-high pressure liquid chromatography system interfaced via a Nanospray Flex nanoelectrospray source (Thermo Fisher). Samples were injected onto a fritted fused silica capillary (30 cm × 75 μm inner diameter with 15 μm tip, CoAnn Technologies) packed with ReprosilPur C18-AQ 1.9 μm particles (Dr. Maisch GmbH). Buffer A consisted of 0.1% formic acid in H_2_O, and buffer B consisted of 0.1% formic acid in 80% acetonitrile. Peptides were separated by an organic gradient from 5% to 35% mobile buffer B over 120 min, followed by an increase to 100% B over 10 min at a flow rate of 300 nL/min. Analytical columns were equilibrated with 3 μL buffer A.

To build a spectral library, samples from each set of biological replicates were pooled and acquired in data-dependent manner. Data-dependent acquisition (DDA) was performed by acquiring a full scan over a m/z range of 375-1025 in the Orbitrap at 120,000 resolving power (@ 200 m/z) with a normalized AGC target of 100%, an RF lens setting of 30%, and an instrument-controlled ion injection time. Dynamic exclusion was set to 30 seconds, with a 10 p.p.m. exclusion width setting. Peptides with charge states 2-6 were selected for MS/MS interrogation using higher energy collisional dissociation (HCD) with a normalized HCD collision energy of 28%, with 3 seconds of MS/MS scans per cycle. Data-independent analysis (DIA) was performed on all individual samples. A full scan was collected at 60,000 resolving power over a scan range of 390-1010 m/z, an instrument controlled AGC target, an RF lens setting of 30%, and an instrument controlled maximum injection time, followed by DIA scans using 8 m/z isolation windows over 400-1000 m/z at a normalized HCD collision energy of 28%.

The Spectronaut algorithm was used to build spectral libraries from DDA data, identify peptides/proteins, and extract intensity information from DIA data. DDA data were searched against the Homo sapiens reference proteome sequences in the UniProt database (one protein sequence per gene, downloaded on October 10, 2019). False discovery rates were estimated using a decoy database strategy. All data were filtered to achieve a false discovery rate of 0.01 for peptide-spectrum matches, peptide identifications, and protein identifications.

An independent mass spectrometry experiment was also performed in collaboration with the lab of Simone Sidoli. Digested peptides were desalted using a 96-well plate filter (Orochem) packed with 1 mg of Oasis HLB C-18 resin (Waters). Briefly, the samples were resuspended in 100 µl of 0.1% TFA and loaded onto the HLB resin, which was previously equilibrated using 100 µl of the same buffer. After washing with 100 µl of 0.1% TFA, the samples were eluted with a buffer containing 70 µl of 60% acetonitrile and 0.1% TFA and then dried in a vacuum centrifuge. Samples were resuspended in 10 µl of 0.1% TFA and loaded onto a Dionex RSLC Ultimate 300 (Thermo Scientific), coupled online with an Orbitrap Fusion Lumos (Thermo Scientific). Chromatographic separation was performed with a two-column system, consisting of a C-18 trap cartridge (300 µm ID, 5 mm length) and a picofrit analytical column (75 µm ID, 25 cm length) packed in-house with reversed-phase Repro-Sil Pur C18-AQ 3 µm resin. Peptides were separated using a 90 min gradient from 4-30% buffer B (buffer A: 0.1% formic acid, buffer B: 80% acetonitrile + 0.1% formic acid) at a flow rate of 300 nl/min. The mass spectrometer was set to acquire spectra in a data-dependent acquisition (DDA) mode. Briefly, the full MS scan was set to 300-1200 m/z in the orbitrap with a resolution of 120,000 (at 200 m/z) and an AGC target of 5x10e5. MS/MS was performed in the ion trap using the top speed mode (2 secs), an AGC target of 1x10e4 and an HCD collision energy of 35. Proteome raw files were searched using Proteome Discoverer software (v2.5, Thermo Scientific) using SEQUEST search engine and the SwissProt human + HIV database. Trypsin was specified as the digestive enzyme with up to 2 missed cleavages allowed. Mass tolerance was set to 10 pm for precursor ions and 0.2 Da for product ions. Peptide and protein false discovery rate was set to 1%.

### MS data analysis

Protein intensities were log_2_ transformed, and log_2_ fold changes in SUMO substrates between the HIV-1 infected versus uninfected samples were calculated as the difference between the average log_2_ abundance of the triplicate HIV-1-infected samples and uninfected samples (Supplemental Table 1 tabs C-F, Supplemental Table 2).

### Endogenous SUMO immunoprecipitation

Immunoprecipitation of SUMO1 or SUMO2/3 conjugated proteins was performed as described^34^ for non-denatured lysates using 50 µL of SUMO1 (Santa Cruz # sc-5308-AC), or SUMO2/3/4 (Santa Cruz # sc-393144 AC) antibody beads per sample in the presence of 15 mM NEM (Sigma #E3876). (SUMO4 is expressed at very low levels relative to SUMO1 and SUMO2/3 in HeLa. Thus, the above antibody was used to immunoprecipitated predominantly SUMO2/3-conjugates). Mouse IgG-Agarose (Sigma # A0919) was used a control. Immunoprecipitated proteins were eluted in 30 µL Laemmli buffer for 5 min. at 95°C on a shaker. Eluates were loaded directly on the gels described below for WB.

### Western blotting

Lysates were prepared using a Millipore Whole Cell Extraction Kit (#2910). Protein content was quantified by Pierce BCA Protein Assay (Thermofisher #23227). Samples were separated on 4-20% Mini-PROTEAN TGX Precast Gels (Bio-Rad #4561096) after heat denaturation in Laemmli buffer. Proteins were transferred to nitrocellulose membranes (Bio-Rad #1620115) using a Transblot-Turbo Transfer System (BioRad). Membranes were blocked in 5% nonfat milk in TBS-T, probed with primary antibodies overnight at 4°C, and incubated with HRP-conjugated secondary antibodies for 1.5h at room temperature. Proteins were detected with SuperSignal West Femto Maximum Sensitivity Substrate (Thermofisher #36095) on an iBright 1500 Imaging System (Thermofisher). Protein levels were quantified using Image Lab (Bio-Rad).

### Antibodies

| Primary antibodies | Catalog # | Dilution |
| --- | --- | --- |
| Monoclonal ANTI-FLAG M2 (Mouse) | Sigma, F3165 | 1:1000 |
| Monoclonal anti-polyHistidine Antibody (Mouse) | Sigma, H1029 | 1:1000 |
| Human SUMO Pan Specific Antibody (Rat) | RD Systems, A-714 | 1:1000 |
| SUMO1 Monoclonal Antibody (21C7) (Mouse) | Invitrogen, 332400 | 1:1000 |
| mAb to Sumo2 + Sumo3 [8A2] (Mouse) | Abcam, AB81371 | 1:1000 |
| Anti-GAPDH mAb (6C5) (Mouse) | Millipore, CB1001 | 1:1000 |
| RanGAP1 (D2T7T) Rabbit mAb | Cell Signaling, 36067S | 1:1000 |
| HNRNPA2B1 pAb (Rabbit) | Proteintech, 14813-1-AP | 1:1000 |
| HNRNPA3 PolyAb (Rabbit) | Proteintech, 25142-1-AP | 1:1000 |
| NACA Polyclonal Antibody (Rabbit) | Invitrogen, 401000 | 3 µg/mL |
| HADHB PolyAb (Rabbit) | Proteintech, 29091-1-AP | 1:1000 |
| mAb to HNRPH2/HNRNPH2 [EPR12171] (Rabbit) | Abcam, AB181171 | 1:1000 |
| HNRNPR PolyAb (Rabbit) | Proteintech, 29980-1-AP | 1:1000 |
| CCAR1 PolyAb (Rabbit) | Proteintech, 25232-1-AP | 1:1000 |
| RPS5 PolyAb (Rabbit) | Proteintech, 16964-1-AP | 1:1000 |
| Vimentin PolyAb (Rabbit) | Proteintech, 10366-1-AP | 1:1000 |
| HERC5 PolyAb (Rabbit) | Proteintech, 22692-1-AP | 1:1000 |
| CHORDC1 PolyAb (Rabbit) | Proteintech, 25525-1-AP | 1:1000 |
| RUVBL2 PolyAb (Rabbit) | Proteintech, 10195-1-AP | 1:1000 |
| UGDH PolyAb (Rabbit) | Proteintech, 13151-1-AP | 1:1000 |
| NOP58 PolyAb (Rabbit) | Proteintech, 14409-1-AP | 1:1000 |
| Histone H1.4 Rabbit mAb (D4J5Q) | Cell Signaling, 41328S | 1:1000 |
| DDX47 PolyAb (Rabbit) | Proteintech, 17795-1-AP | 1:1000 |
| UQCRFS1 PolyAb (Rabbit) | Proteintech, 18443-1-AP | 1:1000 |
| ACTL6A PolyAb (Rabbit) | Proteintech, 10341-1-AP | 1:1000 |
| KHSRP PolyAb (Rabbit) | Proteintech, 55409-1-AP | 1:1000 |
| HNRNPM PolyAb (Rabbit) | Proteintech, 26897-1-AP | 1:1000 |
| HNRNPA1 PolyAb (Rabbit) | Proteintech, 11176-1-AP | 1:1000 |
| SRRM2 mAb (Rabbit) | Abcam, AB317817 | 1:1000 |
| PHF5A PolyAb (Rabbit) | Proteintech, 15554-1-AP | 1:1000 |
| STIP1 PolyAb (Rabbit) | Proteintech, 15218-1-AP | 1:1000 |
| HNRNPA0 PolyAb (Rabbit) | Proteintech, 10848-1-AP | 1:1000 |
| TAF15 PolyAb (Rabbit) | Proteintech, 25521-1-AP | 1:1000 |
| TRA2B PolyAb (Rabbit) | Proteintech, 29627-1-AP | 1:1000 |
| HNRNPU PolyAb (Rabbit) | Proteintech, 16365-1-AP | 1:1000 |
| HNRNPC PolyAb (Rabbit) | Proteintech, 11760-1-AP | 1:1000 |
| SRSF6 PolyAb (Rabbit) | Proteintech, 11772-1-AP | 1:1000 |
| pAb to HIV1 p55 + p24 + p17 (Rabbit) | Abcam, AB63917 | 1:2000 |
| Vpr PolyAb (Rabbit) | Proteintech, 51143-1-AP | 1:1000 |
| mAb to HIV1 Nef (Mouse) | Abcam, AB42355 | 1:1000 |
| Anti-HIV1 Vif antibody (Mouse) | Abcam, ab66643 | 1:1000 |
| Anti-HIV1 Rev antibody (Mouse) | Abcam, ab85529 | 1:1000 |
| Anti-HIV1 Tat antibody (Rabbit) | Abcam, ab43014 | 1:1000 |
| HIV-1 Vpu Antiserum (Rabbit) | Cat # 969 | 1:1000 |
| Anti-GFP antibody (Rabbit) | Abcam, ab290 | 1:1000 |
| hnRNPA2/B1 (B-7) (AC) (used for HNRNPA2B1 IPs) | Santa Cruz, sc-374053<br>AC | 50 μL per IP |
| Rabbit anti-HNRNPA3 (used for HNRNPA3 IPs) | Bethyl, A305-815A-T | 3 μg per IP |
| Normal Rabbit IgG (used as Control for HNRNPA3 IPs) | Cell Signaling, 2729S | 3 μg per IP |

| Secondary antibodies | Catalog # | Dilution |
| --- | --- | --- |
| ECL Anti-Mouse IgG HRP-linked whole antibody | Amersham, NXA931V | 1:10,000 |
| ECL Anti-Rabbit IgG HRP-linked whole antibody | Amersham, NA934V | 1:10,000 |
| EasyBlot anti Rat IgG (HRP) | GeneTex, GTX628474-01 | 1:10,000 |

### Plasmids

All plasmids generated in this study are available upon request. All other plasmids used are commercially available.

### C-terminal SUMO mutants

To generate the 6His-SUMO1^Q92R^ and 6His-SUMO2^Q88R^ expression vectors, gene fragments encoding SUMO1^Q92R^ or SUMO2^Q88R^ were synthesized by GenScript and cloned into pcDNA3-6XHis-human SUMO1 (Addgene #133770) or pcDNA3-6XHis-human SUMO2 (Addgene # 133771), respectively, in place of the WT SUMO sequences. EcoRI and XhoI sites were used for cloning into pcDNA3-6XHis-human SUMO1, and BamHI and XhoI sites were used for cloning into pcDNA3-6XHis-human SUMO2. To generate 3xFlag-SUMO1^Q92R^ and 3xFlag-SUMO2^Q88R^ expression vectors, an insert expressing the 3xFlag tag was synthesized by GenScript and cloned into the above plasmids in place of the 6His tag using KPNI and BamHI sites.

### HIV-1 viral proteins

Plasmids expressing the HIV-1 viral proteins tested in Figure 4D and E are as follows: pALPS vif (Addgene #101328), pALPS vpu (Addgene #101330), pRSV-Rev (Addgene #12253), pcDNA1-Tat (Addgene #138478), and pCMV delta R8.2 (Addgene #12263) for Gag.

### HNRNPA2 lysine mutants

Gene fragments encoding Flag-tagged WT, K108R, K125R, and K305R HNRNPA2 were synthesized by GenScript and cloned into a pCMV3 backbone (SinoBiological) using KPNI and XBAI restriction sites.

### siRNA knockdowns

Cells were plated at 1×10^5^ cells/well of a 6-well plate one day prior to transfection. ON-TARGETplus Human HNRNPA2B1, HNRNPA3, and Non-targeting SMARTpool siRNAs (Dharmacon #s L-011690-01-0005, L-019347-00-0005, and D-001810-10-05) were diluted to stock concentrations of 20 µM. Cells were transfected by dropwise addition of a mixture containing 5 µL of the 20 µM siRNA stocks, 5 µL Lipofectamine RNAiMAX (Invitrogen #13778500) and 490 µL Opti-MEM (Gibco #51985034) at 24 hours post-plating, and again at 48 hours post-plating.

### RNA extraction

Total RNA was extracted using Quick-RNA Miniprep Plus Kit (Zymo #R1058).

### HIV-1 splicing analysis

Splicing quantification was performed as previously described^46^ but using two additional cDNA primers: GTGCTCTTCCGATCTNNNNNNNNNNNNNN uses random bases both for universal priming and as a Unique Molecular Identifier (UMI), GTGCTCTTCCGATCTNNNNNNNNNNTTTYCCACCCCC primes two different sites in the HIV-1 genome at 6257 and 8576. Priming at 6257 identifies IS transcripts and priming at 8576 identifies CS transcripts. cDNA primers were used individually in cDNA reactions and all bead washed cDNAs were used in the subsequent PCRs. The first PCR used ATCTCTCGACGCAGGAC as the forward HIV-1 primer, targeting a region upstream of D1, the major splice donor, and TTCAGACGTGTGCTCTTCCGATCT as the reverse primer, adding additional Illumina platform specific sequences. The second PCR used a nested forward primer combined with Illumina platform sequences GCCTCCCTCGCGCCATCAGAGATGTGTATAAGAGACAGNNNNTGCTG-AAGCGCGCACGGCAAG, and reverse primer TTCAGACGTGTGCTCTTCCGATCT to add additional Illumina sequences, and 5 µl bead washed input from PCR 1. The final PCR used 5 µl bead washed input from PCR 2 and multiplexing primers as in.^46^ Sequencing was done using MiSeq paired-end 300 base reads and demultiplexed using Illumina bcl2fastq pipeline (v.2.20.0). HIV-1 splicing was quantified using a custom pipeline available from the Swanstrom Lab.

### Bioinformatic analysis

GO enrichment analysis was performed with g:Profiler using the Reactome database and Benjamini-Hochberg method to calculate FDR values, where a 5% FDR cutoff was used to select significantly enriched pathways.

### Flow cytometry

Percentages of GFP+ cells were measured using a BD LSRII or LSRFortessa (BD Biosciences). Viable cells were gated using FSC-A and SSC-A plots, and FITC-A and FSC-A plots were used to gate the GFP+ population.

### qPCR

RNA was extracted as described above. cDNA was synthesized using SuperScript III First-Strand Synthesis SuperMix (Invitrogen #11752-050). qPCR was performed on a 7500 Fast Real-Time PCR System using FastStart Universal SYBR Green Master (ROX) (Roche #4913850001). All reactions were performed in triplicate with the following primers: EGFP (forward 5’ AAGGGCGAGGAGCTGTTCACC 3’, reverse 5’ GTGGTCACGAGGGTGGGCCAG 3’), 7SL (forward 5’ATCGGGTGTCCGCACTAAG 3’, reverse 5’ CACCCCTCCTTAGGCAACCT).

### Compounds

10 mM Subasumstat (TAK-981) in 1 mL DMSO was purchased from MedChemExpress (#HY-111789).

### Statistics

Statistical analysis was performed using GraphPad Prism 10 or Microsoft Excel. Error bars indicate mean ± SEM of at least three independent experiments, and significance was determined by unpaired two-tailed t test unless otherwise specified, where p<0.05 was considered statistically significant.

### Data and Software Availability

All additional data and protocols are available upon request. Software used is commercially available. Illustrations were created using BioRender.com.

## ACKNOWLEDGMENTS

This study was funded by Center for Structural Biology of HIV RNA (CRNA) Grant# U54 AI170660, Herbert Irving Comprehensive Cancer Center (HICCC) Grant #P30CA013696, and NIH 1 R01 AI178848. Jeffrey R. Johnson was supported by NIH/NIAID grant R01AI167691. The Sidoli lab gratefully acknowledges for funding the Hevolution Foundation (AFAR), the Einstein-Mount Sinai Diabetes center, and the NIH Office of the Director (S10OD030286).

---

**Supplemental Figure 1.**
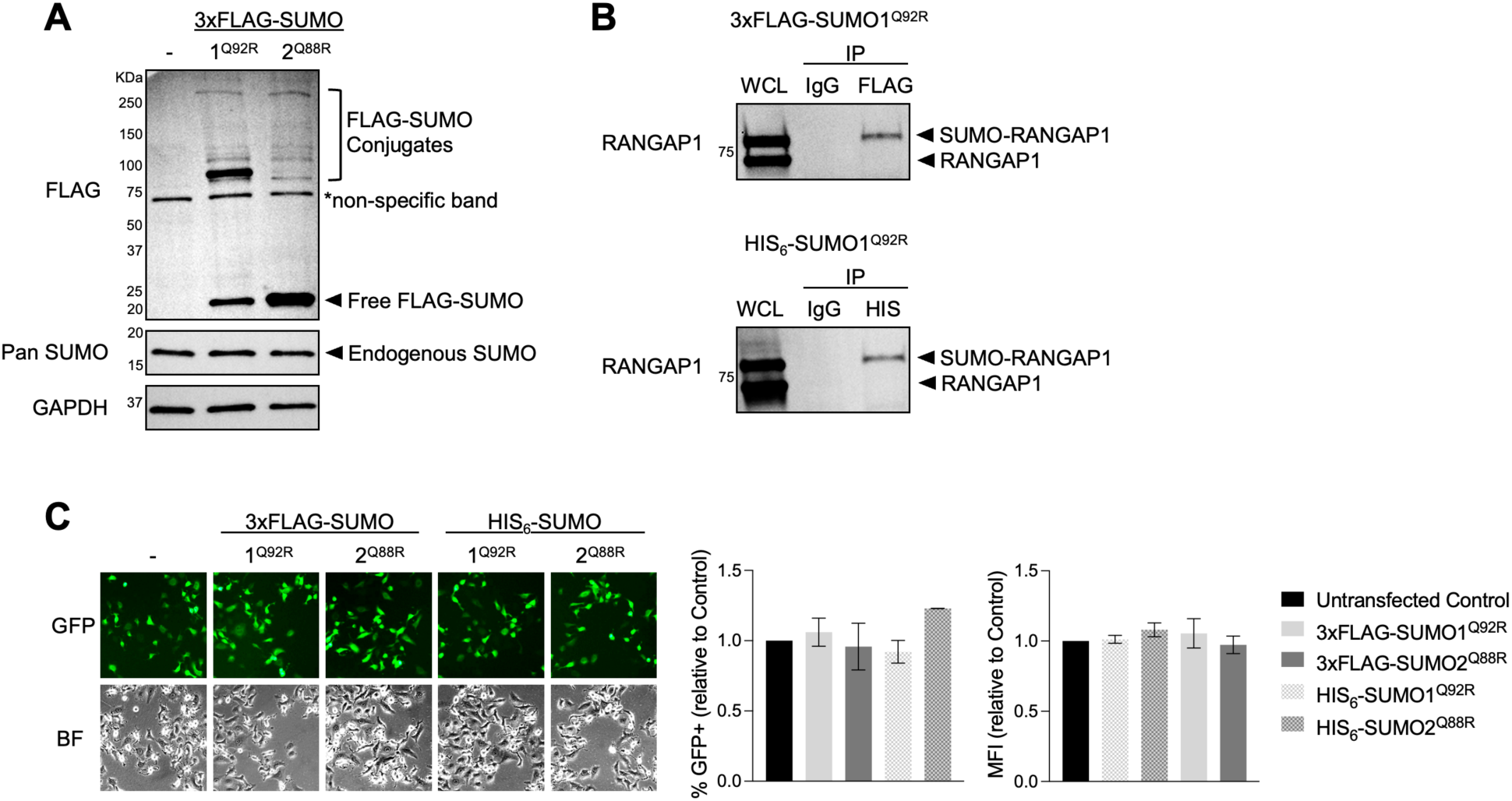
Tagged SUMO mutants are conjugated to endogenous cellular proteins. **(A)** Western blot showing expression of free and conjugated forms of 3xFLAG-SUMO1^Q92R^ and 3xFLAG-SUMO2^Q88R^ in HeLa cells. Expression of endogenous wild-type SUMO is shown for comparison. “-” = Untransfected Control. **(B)** SUMO-modified RanGAP1 is detected following immunoprecipitation with a FLAG or HIS antibody in cells expressing 3xFLAG-SUMO1^Q92R^ or HIS_6_-SUMO1^Q92R^, respectively. Whole cell lysate (WCL) was used as input for IPs. Non-immune IgG = IP control. **(C)** Expression of exogenous SUMO does not interfere with HIV-1 infection. Cell lines were infected with equal volumes of viral supernatants containing VSV-G pseudotyped HIV-1 pNL4-3 deltaENV-EGFP reporter virus. %GFP+ cells and GFP mean fluorescence intensity (MFI) was assessed by flow cytometry at 24 hpi. “-” = Untransfected Control.

**Supplemental Figure 2.**
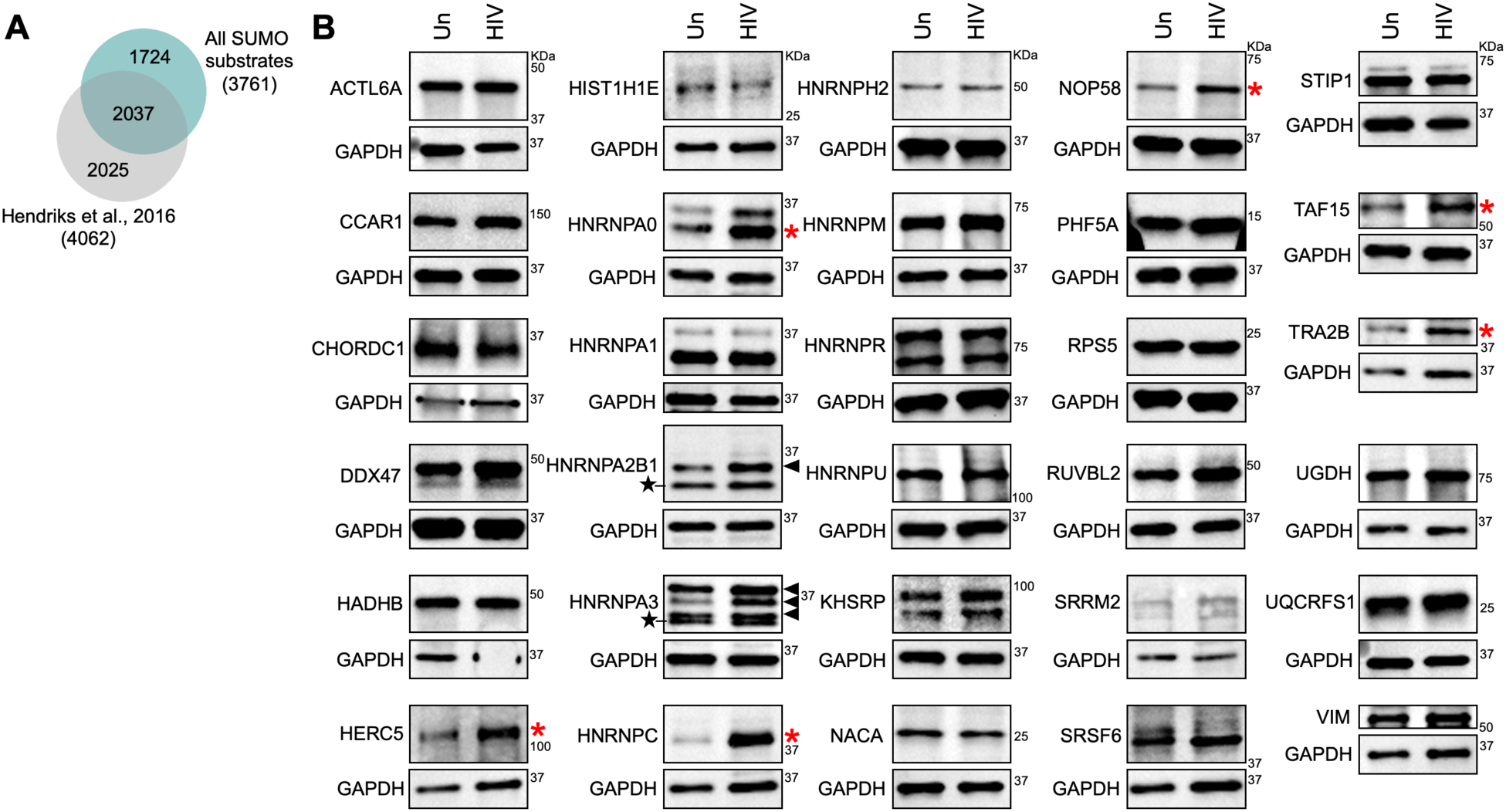
Proteomic screen identifies proteins SUMOylated during HIV-1 infection. **(A)** Overlap between all SUMO substrates identified in this study versus the SUMOylated protein database described in Hendriks et al., 2016. **(B)** Western blots showing expression of select SUMO1 and SUMO2 substrates in HeLa cells left uninfected (“Un”), or infected with HIV-1. Samples were harvested for whole cell lysate preparation at 24 hpi. Arrows indicate SUMO-modified forms of HNRNPA2B1 and HNRNPA3 easily visible on a crude lysate. Stars indicate unmodified forms of these two proteins. Proteins that were increased in total abundance following HIV-1 infection are indicated with red asterisks.

**Supplemental Figure 3.**
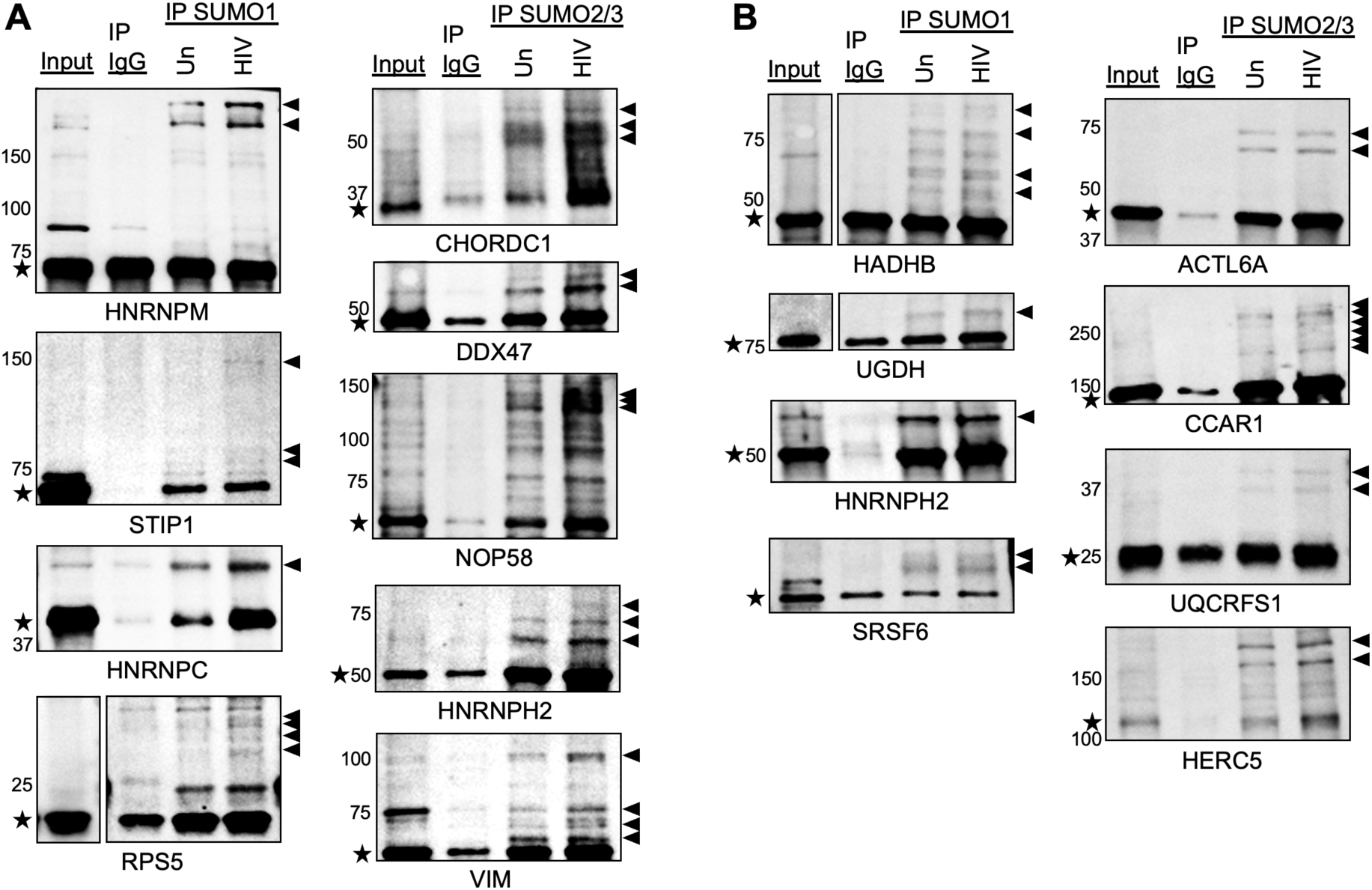
Verification of additional HIV-1 induced SUMO targets. **(A,B)** Western blot analysis of SUMO1- or SUMO2/3-enriched samples from HIV-1-infected cells (”HIV”) versus uninfected controls (“Un”). SUMO-conjugates were immunoprecipitated using anti-SUMO1 or SUMO2/3 antibodies at 24 hpi as described in Figure 2, and samples were analyzed by western blot for the additional candidate substrates indicated below each blot. Stars indicate unmodified proteins. Arrows indicate SUMO-conjugated proteins. Crude lysate from HIV-1-infected cells was used as input for IPs. Input is 5% of protein loaded in IPs. Non-immune IgG = IP control.

**Supplemental Figure 4.**
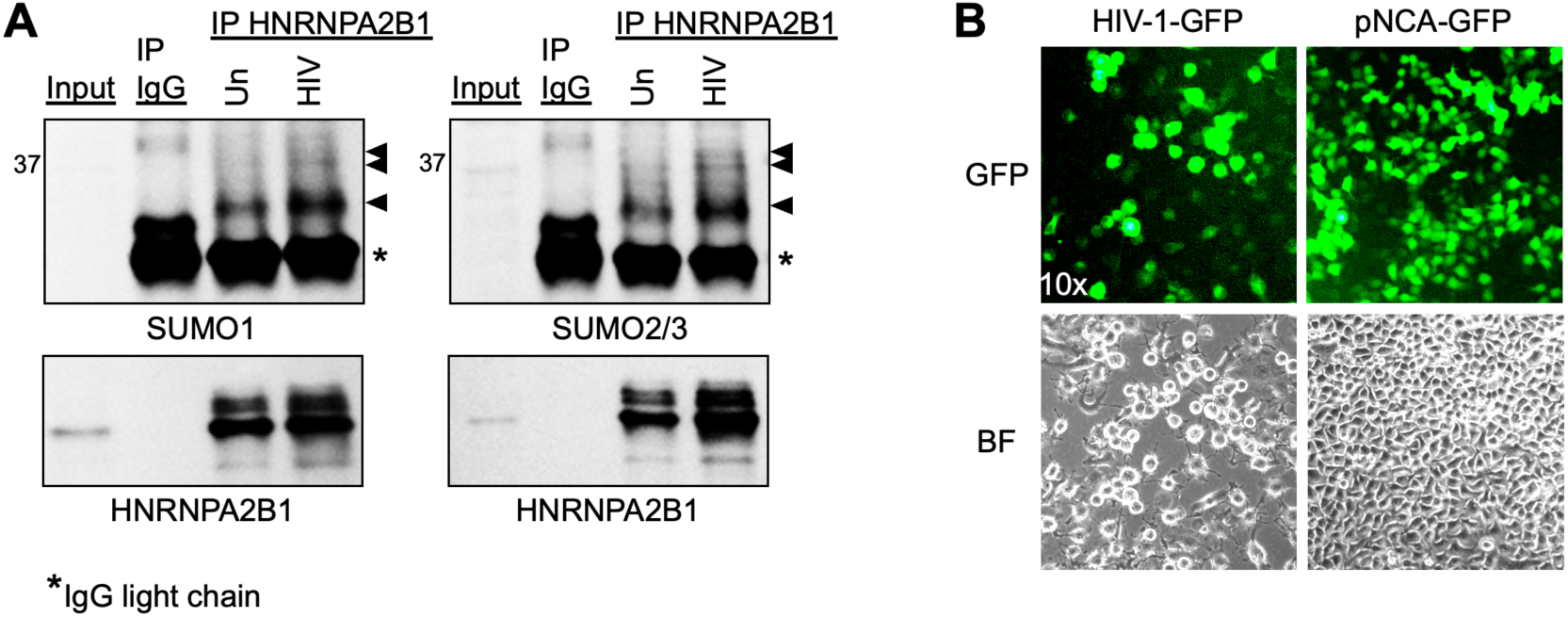
Validation of HIV-1-induced hnRNPA2B1 SUMOylation. **(A)** Western blot analysis of HNRNPA2B1-enriched samples for SUMO1 and SUMO2/3 conjugation. HNRNPA2B1 was immunoprecipated from uninfected control (“Un”) versus HIV-1-infected cells at 24 hpi., and samples were analyzed by western blot with anti-SUMO1 or SUMO2/3 antibodies. Arrows indicate SUMO-conjugated proteins. Asterisks indicate IgG light chain. Crude lysate from HIV-1-infected cells was used as input for IPs, where input was 5% of protein loaded in IPs. Non-immune IgG = IP control. **(B)** Representative images showing GFP expression in HeLa cells infected with VSV-G pseudotyped HIV-1 GFP versus MMLV reporter pNCA-GFP. Cells are shown at 48 hpi.

**Supplemental Figure 5.**
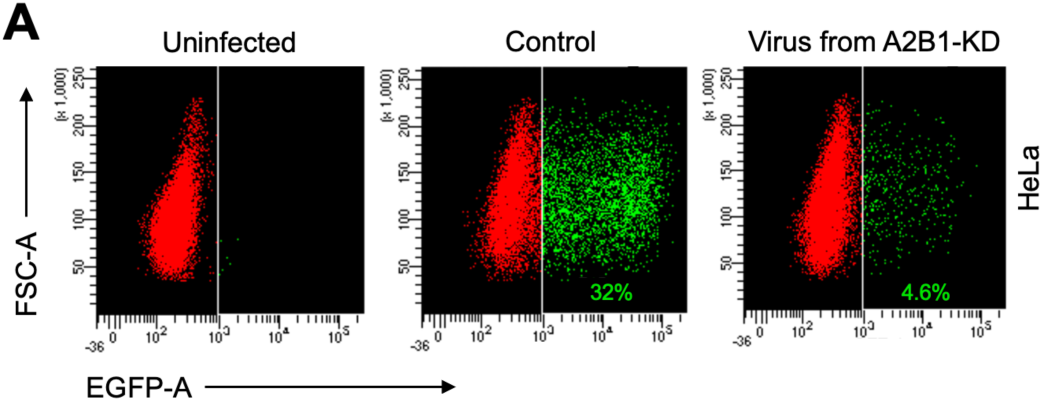
Effect of HNRNP A2B1 and A3 knockdown on HIV-1 splicing and virion production. **(A)** Flow cytometry measuring GFP expression in HeLa cells infected with HIV-1-GFP virus produced by Control versus A2B1-KD 293T cells. GFP expression was measured at 24 hpi. Shown are representative images of three independent experiments.

**Supplemental Figure 6.**
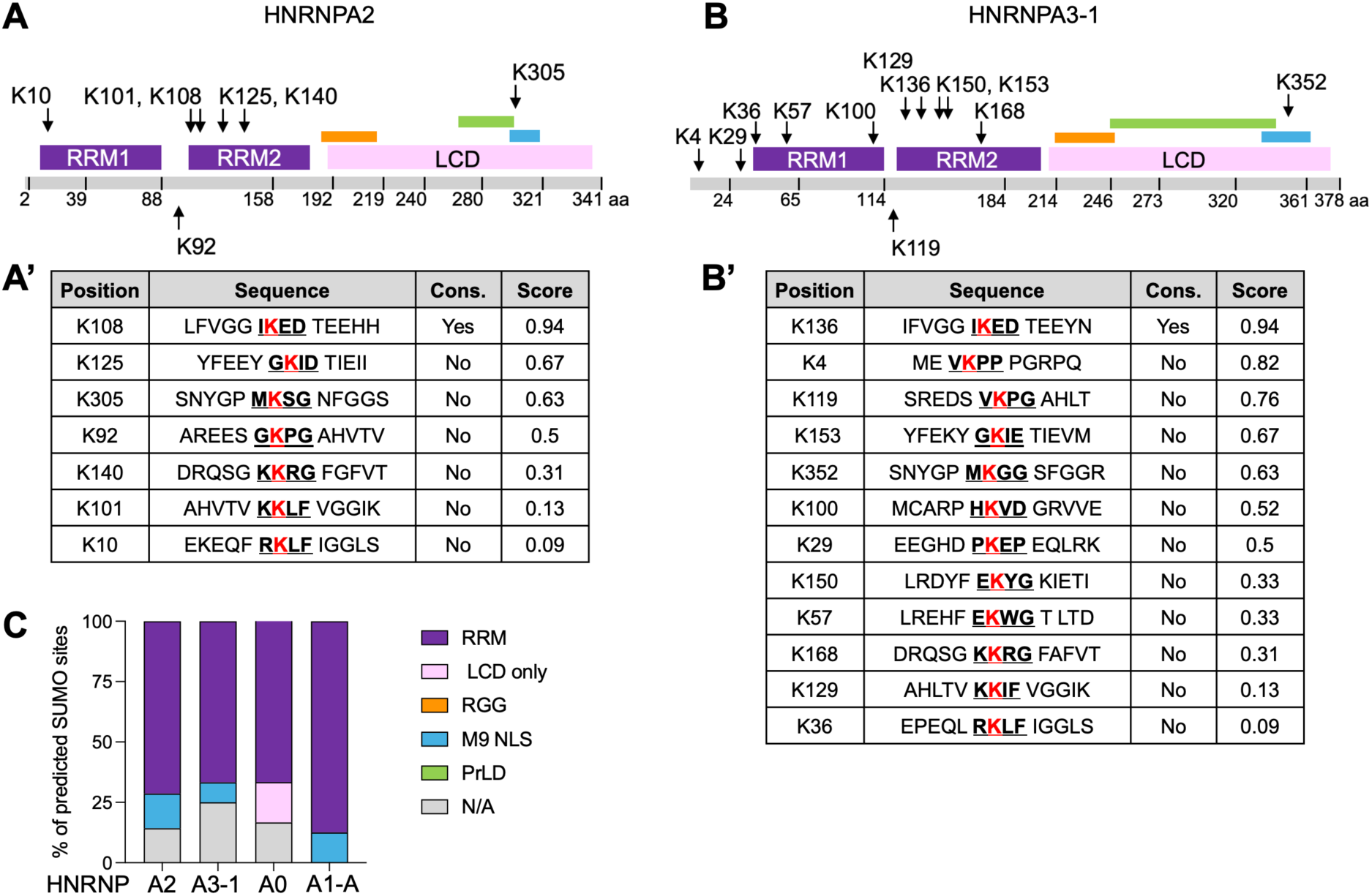
HNRNPA/B SUMOylation sites suggest a role in modulating RNA-binding. **(A,B)** Schematic of the HNRNPA2 and HNRNPA3-1 isoforms, the most abundant splice isoforms of A2B1 and A3, respectively. Total amino acid sequence is shown in light gray, with numbers denoting each exon. Purple denotes the RNA Recognition Motifs (RRM) 1 and 2, pink denotes the glycine-rich, low complexity domain (LCD). Contained largely within the LCD, are the arginine-glycine-glycine (RGG) box (orange), M9 nuclear localization sequence (blue), and prion-like domain (PrLD) (green). The location of predicted SUMOylated lysine (K) residues listed in parts A’ and B’ are indicated. **(A’, B’)** Table of predicted SUMOylated K residues in the HNRNPA2 and HNRNPA3-1 isoforms. “Cons.” indicates the presence or absence of the SUMOylation consensus motif. Score indicates the probability of a site being SUMOylated, where a score of 1 indicates highest probability. **(C)** Stacked bar graph showing the percentage of predicted SUMOylation sites located in the RRM, LCD only, RGG, M9 NLS, or PrLD of HNRNPs A2, A3-1, A0, and A1-A. SUMO sites not located in any of these domains are listed in “N/A.

